# Complement Biosensors Identify a Classical Pathway Stimulus in Complement-Mediated Hemolytic Uremic Syndrome

**DOI:** 10.1101/2024.05.29.596475

**Authors:** Michael A. Cole, Nikhil Ranjan, Gloria F. Gerber, Xiang-Zuo Pan, Daniel Flores-Guerrero, Shruti Chaturvedi, C. John Sperati, Keith R. McCrae, Robert A. Brodsky

## Abstract

Complement-mediated hemolytic uremic syndrome (CM-HUS) is a thrombotic microangiopathy characterized by germline variants or acquired antibodies to complement proteins and regulators. Building upon our prior experience with the modified Ham (mHam) assay for ex vivo diagnosis of complementopathies, we have developed an array of cell-based complement “biosensors’’ by selective removal of complement regulatory proteins (CD55 and CD59, CD46, or a combination thereof) in an autonomously bioluminescent HEK293 cell line. These biosensors can be used as a sensitive method for diagnosing CM-HUS and monitoring therapeutic complement blockade. Using specific complement pathway inhibitors, this model identifies IgM-driven classical pathway stimulus during both acute disease and in many patients during clinical remission. This provides a potential explanation for ~50% of CM-HUS patients who lack an alternative pathway “driving” variant and suggests at least a subset of CM-HUS is characterized by a breakdown of IgM immunologic tolerance.

**Key Points:**

1. CM-HUS has a CP stimulus driven by polyreactive IgM, addressing the mystery of why 40% of CM-HUS lack complement specific variants
2. Complement biosensors and the bioluminescent mHam can be used to aid in diagnosis of CM-HUS and monitor complement inhibitor therapy

## Introduction

Complement-mediated hemolytic uremic syndrome (CM-HUS) is a subset of thrombotic microangiopathy (TMA). CM-HUS remains a clinical diagnosis based on exclusion of other TMA syndromes; there is no diagnostic assay. Prompt diagnosis of CM-HUS is critical as complement inhibition with C5 inhibitors is highly effective therapy^1^.

Complement-mediated HUS is thought to be a disorder of the alternative pathway (AP) of complement as 50-60% of patients harbor a germline variant or acquired autoantibody to a protein central to this pathway (eg, C3, FH, FI, MCP/CD46, CFB)^2,3^. Mechanistic studies show these pathogenic variants play a direct role in either cell surface expression, gain of function/resistance to regulation, cofactor activity, or decay accelerator activity^4–6^. However, these variants are neither necessary nor sufficient; 40-50% of patients do not carry any variant or autoantibody, and disease penetrance is only 20-50% in those with a known pathogenic variant^7,8^. Thus, there is an unmet need for assays and/or biomarkers not only for diagnosis, but also to mechanistically explain disease in patients without variants and the low penetrance in those with known familial pathogenic variants.

Two principal techniques for *in vitro* diagnosis have been proposed, including the modified Ham assay (mHam)^9,10^ and HMEC-1 staining for C5b-9^11–14^, but neither is widely adopted. Both assays rely upon testing patient serum for abnormal cell-deposited complement activity. In the case of the mHam, TF-1 erythroblasts deficient in glycosylphosphatidylinositol (GPI)-linked complement regulators (DAF/CD55 and CD59) are used in an absorbance-based cell viability assay to assess for increased complement activity leading to cell death^9^. In most versions of the HMEC-1 C5b-9 assay, wild type HMEC-1 endothelial cells are stimulated with ADP, incubated with patient serum, and then assessed by indirect immunofluorescence microscopy for increased C5b-9 deposition^12^.

Our experience with the mHam assay suggested an additional stimulus beyond the AP was required for complement activation. First, complement-mediated cell killing was typically greater in calcium and magnesium-containing buffer permissive of all pathways (classical, lectin, and alternative) than in an AP-specific buffer (MgEGTA)^9^. Second, patients with known pathogenic variants in membrane bound (versus soluble) complement regulators (e.g, CD46) were still positive when testing their serum in either the mHam or HMEC-1 C5b-9 assays^13,15,16^. Finally, both assays, as well as sC5b-9, remain positive or elevated in a sizable proportion of patients without an identifiable AP variant, *even* in hematologic remission^11,13,17^. The initiation of the AP is hydrolysis of the thioester of C3 which occurs at a slow, fixed rate according to first order kinetics^18,19^. Although enhanced tickover through contact activation or heme-induced activation has been proposed as a contributing mechanism, this would not explain ongoing complement activation in a CM-HUS patient in hematologic remission^18,20–22^. Taken together, these data suggest an additional mechanism of complement activation contributes to disease pathogenesis.

To address these observations, we used an autonomously bioluminescent HEK293 cell line which allows for real time measurement of cellular metabolic health resulting from complement activation and obviates the need for washes or reagent addition^23^. As compared to the TF-1 cell line used in the mHam, HEK293 cells lack complement receptor 1 (CR1), which strongly regulates both classical pathway (CP) and AP activity^24^ on cells of hematopoietic origin and renal podocytes^25^. Therefore, HEK293 cells more closely approximates complement activation of relevance to the endothelium in CM-HUS. By removal of different membrane-bound complement regulatory proteins, we established three autonomously bioluminescent HEK293 biosensors with different susceptibilities to complement activation: an isolated MCP/*CD46* knockout (*CD46*KO), a *PIGA* knockout (*PIGA*KO) which results in deficiency of DAF/CD55 and CD59, and a double *PIGA* and *CD46* knockout line (DKO) deficient in all three regulators. Residual membrane level complement protection in the DKO is provided only from Factor H (FH) or related proteins.

Here, we use these biosensors to probe the pathogenesis of CM-HUS and show the presence of abnormal CP activity driven by IgM.

## Methods

### Patient Cohort and Definitions

We utilized 41 biobanked sera specimens from the Johns Hopkins Complement Associated Disease Registry with a diagnosis of CM-HUS or TTP (Table 1). TTP was diagnosed by ADAMTS13 activity <10% and the diagnosis of CM-HUS was determined by a consulting hematologist. Samples were defined as acute if they were collected within 14 days of initial manifestations, 2) had not yet received donor plasma, C5 inhibitory therapy or caplacizumab, and 3) had not yet achieved hematologic remission (defined as Hgb >11 g/dL or recovery to baseline, LDH within normal limits and platelet count >150×10^3^/μL). Patients with renal-limited TMA were excluded. Two of 6 (33%) acute CM-HUS samples and 6 of 15 (40%) remission samples had a known pathogenic variant or variant of uncertain significance (VUS) in a complement regulatory protein or factor H autoantibody (FHAA). Fourteen CM-HUS samples were included from patients on C5 inhibitors (either eculizumab or ravulizumab). Additionally, we included sera from 19 non-pregnant, healthy adults and 6 acute TTP patients.

**Table 1.** Cohort characteristics.

| Patient identification | Age at diagnosis (y)/ Sex | Sample collected/ Follow-up (y)<sup>a</sup> | Variant<sup>b</sup> | Overall classification | Trigger | Therapy<sup>c</sup> | Clinical outcome |
| --- | --- | --- | --- | --- | --- | --- | --- |
| **Acute** |  |  |  |  |  |  |  |
| CM01 | 27/F | 0/4 | NA | NA | 3<sup>rd</sup> pregnancy | Short-term eculizumab | Normal kidney function |
| CM02 | 14/F | 0/4 | CD46 c.286+2T>G splice site | Likely pathogenic | Influenza A | Supportive care only | Normal kidney function |
| CM03 | 9/M | 0/4 | +FHAA, homozygous del of *CFHR1*, compound heterozygous *CFHR3-CFHR1* and *CHFR1-CHFR4* deletion | Likely pathogenic | Influenza B | Long-term eculizumab | Normal kidney function |
| CM04 | 23/F | 0/5 | Homozygous del *CHFR3-CHFR1*, no FHAA | VUS | NA | Short-term eculizumab | Deceased (unrelated cause) |
| CM05 | 1/M | 0/3 | NMD | NA | *S. pneumoniae* meningitis | Supportive care | Normal kidney function |
| CM06 | 59/F | 0/<1 | NA | NA | NA | On eculizumab | ESKD |
| **Remission** |  |  |  |  |  |  |  |
| CM07 | 46/F | 4/11 | Homozygous del of *CFHR1-CHFR3* and heterozygous *Factor H* p.Gln950His | Likely pathogenic vs VUS | Kidney stone | Short-term eculizumab | CKD1 |
| CM08 | 22/F | 3/5 | NMD | NA | Possible infection | Short-term eculizumab | Normal kidney function |
| CM09 | 22/F | 4/9 | NMD | NA | Flu-like illness | Short-term eculizumab | CKD3 |
| CM10 | 31/F | 1/4 | NMD | NA | pregnancy | PLEX/Short-term eculizumab | ESKD |
| CM11 | 26/F | 1/2 | NMD | NA | pregnancy | short-term eculizumab | CKD1 |
| CM12 | 24/F | 1/1 | Heterozygous *CD46* c.104G>A, p.Cys35Tyr | Likely pathogenic | COVID-19 | Supportive care | Normal kidney function |
| CM13 | 24/M | 2/8 | *ADAMTS13* -c.3677C>T, p.Thr1226Ile | VUS | Possible pneumonia | Short-term eculizumab | CKD4 |
| CM14 | 31/F | 3/11 | *THBD* (heterozygous c.1456G>T, p.Asp486Tyr), *CFHR5* (heterozygous c.1357C>G, p.Pro453Ala) | Likely pathogenic | pregnancy | PLEX and rituximab | Normal kidney function |
| CM15 | 55/M | 1/2 | NA | NA | EtOH pancreatitis | Short-term eculizumab | Normal kidney function |
| CM16 | 37/F | 1/8 | *THBD* het *c.127G>A*, p.Ala43Thr and het del *CFHR3-CFHR1* with novel fusion protein of the 1st 19 SCR of CFH and SCR 5 of CFHR1 | Likely pathogenic | Possibly IBD related | Long-term ravulizumab | Normal kidney function |
| CM17 | 51/M | 1/8 | het *CD46* p.Ser274Tyrfs*, het *CFH* p.Arg1074Pro, het del *CFHR3-CFHR1* | Likely pathogenic | Unknown | Short-term eculizumab | Normal kidney function |
| CM18 | 24/F | 1/9 | NMD | NA | Dental abscess | Supportive | Normal kidney function |
| CM19 | 22/F | 2/11 | NMD | NA | Unknown | Short-term | ESKD s/p transplant |
| CM20 | 37/F | 1/2 | NMD | NA | Pregnancy | PLEX only | CKD3 |
| CM21 | 2/M | 27/28 | Het C3 c.Arg592Trp | Likely pathogenic | Unknown | Long-term ravulizumab | ESKD s/p transplant (x3) |

NMD = No mutation detected. NA = not performed or not available. PLEX = plasma exchange. <sup>a</sup>Sample collection indicates time from diagnosis until sample was collected in years. Length of follow-up indicates time from diagnosis in years until present. <sup>b</sup>In only two cases of CFHR1/CFHR3 deletion was FHAA specifically tested (CM03/CM04). Given other cases are remission samples, they were not analyzed for FHAA but classified as likely pathogenic vs VUS based upon the homozygous deletion and known increased frequency in CM-HUS. <sup>c</sup>Short-term defined as a minimum of 5 doses or up to two years and around the time of kidney transplant in some cases.

### Biosensor Creation

LiveLight HEK293 cell line (490 BioTech) was utilized to allow for real time measurement of metabolic health^23^. A *PIGA* knockout cell line (*PIGA*KO) devoid of all GPI-anchored proteins, including complement regulators CD55 and CD59, was established by proaerolysin selection as previously described^10^.

CRISPR/Cas9 was used to establish an isolated MCP/CD46 knockout cell line (*CD46*KO) as well as a double knockout cell line (DKO) devoid of all three membrane bound complement regulators, CD55, CD59, and CD46.

### Bioluminescent mHam Assay

Forty-thousand cells were harvested with trypsin/EDTA and plated as triplicates in gelatin veronal buffer (GVB^++^, 80 μL). Next, serum (20 μL) was added after appropriate treatment (heat inactivation, DTT treatment, IdeS cleavage, or inhibitor addition as described in supplemental methods). Relative luminescence was monitored by serial luminescence measurements (every 5 min at 37 °C) in a BMG Clariostar luminometer. Results were plotted in GraphPad Prism.

### Pre-sensitization

CM-HUS or HC sera was heat inactivated and then incubated for 45 min at 37 °C with *PIGA*KO or DKO cells. Cells were washed twice and resuspended in GVB^++^. These pre-sensitized cells are then utilized in the bioluminescent mHam as described above with HC mix serum (specifically selected for low activity on the DKO cell line).

### Flow Cytometry

Healthy control or CM-HUS serum was incubated for 30 min at 37 °C with wild type TF-1, HEK293, *CD46*KO, and *PIGA*KO cell lines in the presence of eculizumab alone, eculizumab + sutimlimab, or eculizumab + factor D inhibitor (ACH-5548). Heat-inactivated sera was used as a negative control. Cells were washed in Hanks balanced salt solution with 1% BSA, stained for C3c or C4d, and analyzed by flow cytometry as described in the supplemental methods. IgG and IgM flow cytometry was similarly performed after incubation of heat-inactivated healthy control or CM-HUS sera for 45 min at 37 °C.

### Statistics and Illustration Preparation

Descriptive statistics and statistical analysis were performed in GraphPad Prism. Graphical illustrations were created with BioRender and Adobe Illustrator.

A complete description of experimental details can be found in the supplemental materials.

## Results

### Biosensors detect range of normal complement activity among healthy controls

In the bioluminescent mHam, cells are incubated with sera for 2.5 hours and a real time tracing of autonomous bioluminescence is compared to heat-inactivated sera (Fig. 1A-E). Heating serum is a well-established method for inactivating complement^26^, and cell luminescence correlates with metabolic health and cell viability^23,27^. Thus, by comparing the luminescent signal of untreated and heat-inactivated sera from the same patient, the effects of complement-mediated cell cytotoxicity are measured. As maximal effect is typically observed at ~1 h, this time point was selected to report a “percent relative luminescence,” where 100% relative luminescence represents no significant metabolic effect of complement and zero percent indicates total loss of the cell population. Using healthy control sera, the relative luminescence at 1 h decreased as complement regulators were removed (Fig 1B-F): wild type (WT) cells 90.3 (95% CI, 86.9-93.8), *CD46*KO cells 78.8 (95% CI, 67.9-89.7), *PIGA*KO cells 47.2 (95% CI 34.4-59.9), and DKO 20.9 (95% CI 7.9-33.9) (Fig. 1F). One healthy control was excluded given significant killing on WT cells with further studies ongoing to determine the etiology (Fig. 1F). A relative 1 h luminescence of ≤12% for the *PIGA*KO and ≤62% for the *CD46*KO line was used to establish a “positive” threshold for abnormal complement activity, corresponding to the lower 85^th^ percentile on both cell lines.

**Fig. 1.**
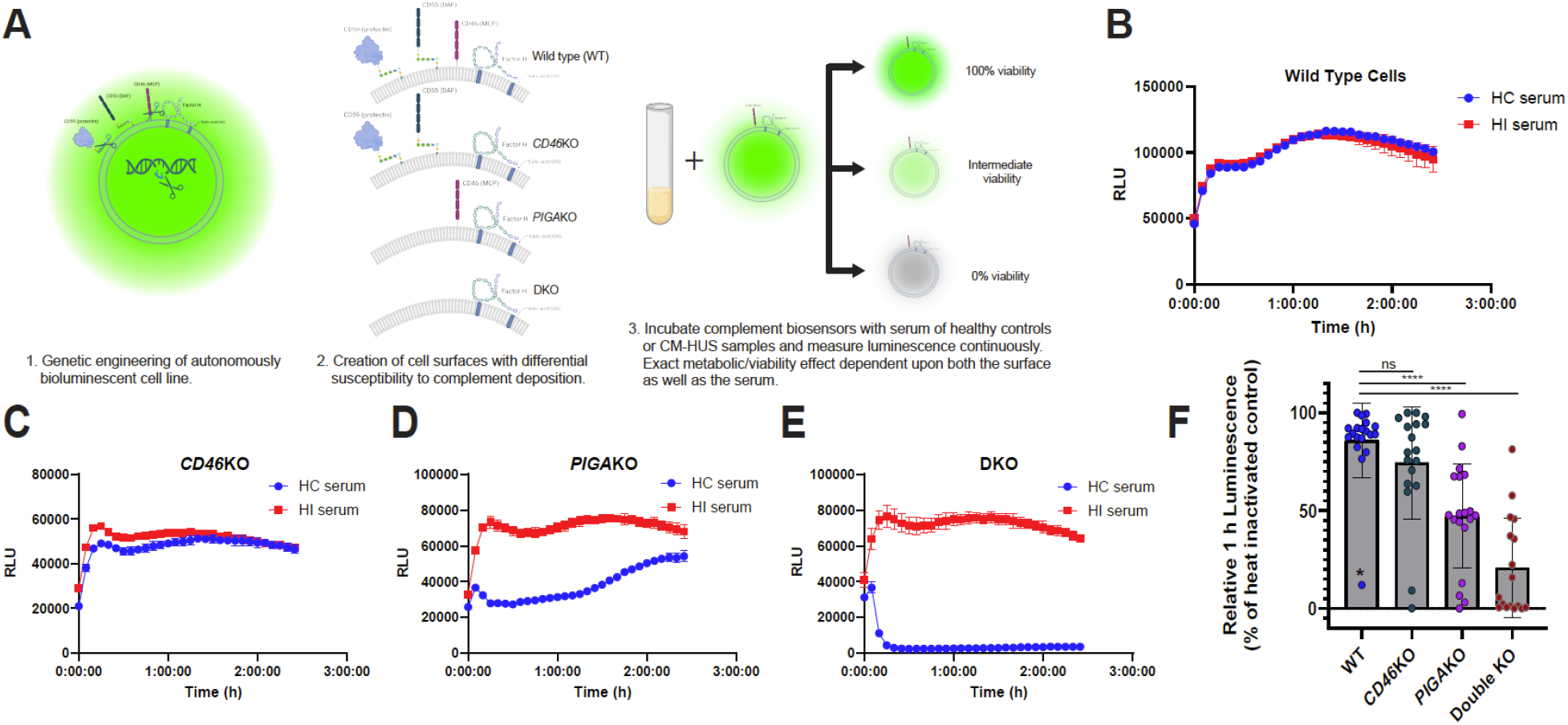
Complement biosensors identify a range of “normal” complement activation in healthy control sera. **A**, Overview of creation of biosensors with different susceptibilities to complement and description of the assay. Created with biorender.com. **B-E**, Livelight WT HEK293 cells (**B**), *CD46*KO cells (**C**), *PIGA*KO cells (**D**), Double *CD46* and *PIGA* KO (DKO) cells (**E**) incubated with healthy control (HC) or heat-inactivated sera from the same HC. Relative luminescence measured every 5 minutes for 2.5 h. RLU = relative luminescence units. Data plotted as mean +/- SD for each triplicate. **F**, Summary of relative luminescence at t = 1 h for healthy controls. n= 19 for WT, *CD46*KO, *PIGA*KO; n=18 for DKO. (*) identifies outlier by ROUT analysis (Q=1%). *P* values were calculated using one-way ANOVA for Dunnett’s multiple-comparisons test. ns, not significant, *p < 0.05, **p < 0.01, ***p < 0.001, ****p < 0.0001.

### Bioluminescent mHam on *PIGA*KO and *CD46*KO biosensors differentiates acute CM-HUS from TTP and healthy control sera

We next sought to characterize the activity in thrombotic microangiopathy samples, including CM-HUS (acute or remission) and acute TTP. Both acute and remission CM-HUS samples are significantly different than healthy controls or acute TTP, and all acute CM-HUS have significant activity on both the *PIGA*KO and *CD46*KO biosensors (Fig. 2A,B,D,E). In remission, 54% (7/13) of samples remain positive on the CD46KO and 57% (8/14) on the PIGAKO (Fig. 2B,E). Among patients with a complement variant or FHAA, 60% (3/5) were positive on *CD46*KO as compared to 50% (4/8) without.

**Fig. 2.**
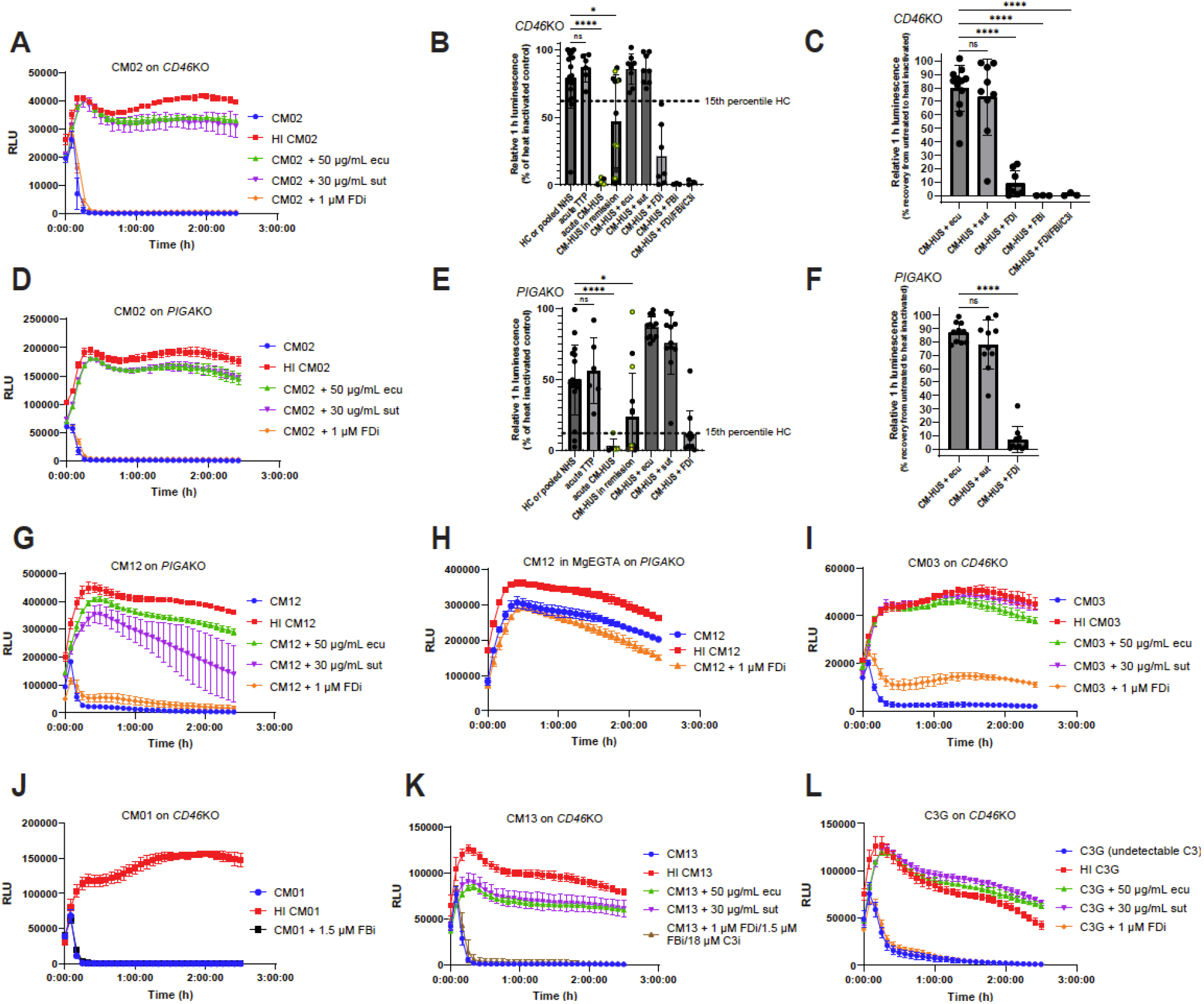
Example CM-HUS tracings and summary of relative luminescence and blocking. **A,D**, CM02 sera on *CD46*KO cells (**A**) or *PIGA*KO cells (**D**) in GVB++ with or without heat inactivation (HI), eculizumab (ecu, 50 μg/mL), sutimlimab (sut, 30 μg/mL), or ACH-5548 (FDi). **B,E**, Summary of relative 1 hour luminescence of healthy controls (HC), CM-HUS (acute or remission) with or without the addition of eculizumab, sutimlimab, ACH-4471/ACH-5548 (FDi, 1 μM), iptacopan (FBi, 1.5 μM), or “triple AP blockade” with FDi (1 μM), FBi (1.5 μM), and compstatin (18 μM) on *CD46*KO (**B**) or *PIGA*KO (**E**). *PIGA*KO: HC n=18, acute TTP n=6, acute CM-HUS n=5, remission n=14, CM-HUS + ecu n=12, CM-HUS + sut n=11, CM-HUS + FDi n =11. *CD46*KO: HC n=18, acute TTP n=6, acute CM-HUS n=5, remission CM-HUS n =13, CM-HUS + ecu n=8, CM-HUS + sut n=8, CM-HUS + FDi n =7, CM-HUS FBi n=3, CM-HUS triple blockade n=3. Graphed as mean +/- SD. Yellow fill indicates sample with pathogenic variant or VUS in complement regulatory protein. Dashed line represents the 15th percentile of healthy control. **C,F**, Relative 1 h luminescence recovery after addition of inhibitor calculated as [(luminescence inhibitor treated serum-luminescence untreated serum)/(luminescence heat inactivated serum-luminescence untreated serum)] at t=1 h. *PIGA*KO: n=10 for each condition, *CD46*KO: CM-HUS + ecu n=12, CM-HUS + sut n=9, CM-HUS + FDi n=8, CM-HUS +FBi n=3, triple blockade n=3. **B,C,E,F**, *P* values were calculated using one-way ANOVA for Dunnett’s multiple-comparisons test. **G,H**, AP only buffer inhibits complement activity. CM12 sera on *PIGA*KO in GVB++ (**G**) or MgEGTA (AP only buffer with 13 mM Mg2+) (**H**) with or without inhibitors as indicated in legend. **I**, Example tracing of acute CM-HUS CM03 (Factor H autoantibody) on *CD46*KO with or without inhibitors as indicated in legend. **J**, Example tracing of remission CM-HUS no mutation CM01 on *CD46*KO with addition of iptacopan (FBi, 1.5 μM). **K**, Example tracing of CM13 on *CD46*KO with addition of “triple AP blockade” including with FDi (1 μM), FBi (1.5 μM), and compstatin (18 μM). **l**. Example tracing of C3 glomerulopathy (C3G) with VUS in *CD46* on *CD46*KO. Activity persists despite C3 being below limit of detection at time sample was collected (<15 mg/dL). **A,D,G,H,I,J,K,L**, RLU = relative luminescence units. Data plotted as mean +/- SD for each triplicate. ns, not significant, *p < 0.05, **p < 0.01, ***p < 0.001, ****p < 0.0001.

### Use of classical pathway C1s inhibitor sutimlimab but not AP inhibitors mitigates cell-directed complement activation in CM-HUS sera

To investigate the contribution of each complement pathway, we utilized a selection of complement inhibitors: AP inhibitors-factor D inhibitors (FDi) ACH-4471 and ACH-5548, factor B inhibitor (FBi) iptacopan, C3 inhibitor compstatin; CP inhibitor-C1s inhibitor sutimlimab; terminal pathway-C5 inhibitor eculizumab. Activity of these inhibitors was confirmed in the AP or CP Wieslab assays (Fig. S1). The terminal pathway inhibitor eculizumab rescued the bioluminescent signal to a mean of 80% on *CD46*KO and 87% on *PIGA*KO (Fig. 2C,F), but unexpectedly, AP inhibitors offered minimal protection from complement-mediated cell death. With FDi, there was mean recovery of 8.8% on *CD46*KO and 6.8% on *PIGA*KO (Fig. 2A-G,I). This was confirmed with a FBi on a limited number of samples (Fig. 2B,C,J). In contrast, the CP inhibitor sutimlimab offered significant protection and was comparable to eculizumab with mean recovery of 73% on *CD46*KO and 78% on *PIGA*KO (Fig. 2C,F). The sutimlimab findings were confirmed by using MgEGTA, a buffer permissive of only AP, which also blocked the serum cytolytic activity (Fig. 2G,H). These effects were consistently observed in both acute and remission samples, regardless of the presence of an AP variant (compare Fig. 2A,B,E,I,K). Importantly, investigations on a sample with known factor H autoantibody level of 57,104 arbitrary units (<200 AU normal), CM03, showed the highest degree of blockade with a factor D inhibitor (27.7%) (Fig. 2I), but this was still less than blockade with sutimlimab (98.6%). Additionally, healthy control sera in MgEGTA on the DKO line shows that alternative pathway can be detected and appropriately blocked by FDi but not by sutimlimab (Fig. S2). The pattern of the curves in MgEGTA buffer (delayed as opposed to early cell kill) also highlight that as expected alternative pathway plays a more important role in *amplification* rather than as an initiating stimulus^22,28^. Furthermore, pathologic complement activation in sera from patients with variants in membrane bound regulators (eg CD46) was blocked by sutimlimab but not AP inhibitors in both acute and remission samples (Fig. 2A,D,G). “Triple” AP blockade targeting FB, FD, and C3 did not offer significant protection (Fig. 2B,C,K). Finally, the sera of a patient with C3 glomerulopathy (Table S2) with undetectable C3 (<15 mg/dL) showed reduced bioluminescence comparable to CM-HUS (Fig. 2L). Given that complement activity persists with either C3 depletion or triple AP blockade, the “C3 bypass mechanism” may lead to membrane attack complex (MAC) deposition. This supports observations of others that in the presence of strong CP stimulus on membrane surfaces, there is a more limited role for AP amplification^29,30^. These findings suggest that regardless of AP dysregulation, there is a CP stimulus in CM-HUS. Highlighting the membrane-directed nature of the CP stimulus, heat aggregated IgG, a well-documented fluid phase CP activator, *protects* cells from death in a dose-dependent manner (Fig. S3). Furthermore, this exemplifies the important differences between fluid phase analysis and assays of membrane-directed activation^31,32^.

### Classical pathway stimulus drives C4d deposition in CM-HUS even on wild type HEK293 and TF-1 cells, and MCP/CD46 as opposed to CD55 is a more significant regulator of that stimulus

Of the membrane associated complement regulators, only variants in MCP (CD46) and Factor H are associated with CM-HUS^6,33^. In addition to AP regulation, these proteins are C4b binding proteins and regulate CP activity with variable efficiency depending upon the model system and degree of sensitization^34–38^. Aside from CR1, they are the only membrane associated regulators with cofactor activity to control CP-stimulated C4b deposition on the cell surface ^25,36^. DAF (CD55) regulates deposition of both CP and AP convertases but possesses only decay accelerator activity^6,39^.

To study the role of CD46 and CD55 in CM-HUS, we performed flow cytometry to assess C3c and C4d deposition in CM-HUS compared to healthy control sera (Fig. 3). Eculizumab was utilized to preserve cell viability. Given CD59 inhibits only MAC insertion (ie downstream of C3c and C4d deposition), differential deposition on WT, *PIGA*KO, and *CD46*KO will be driven by either the lack of DAF/CD55 (for *PIGA*KO line) or MCP/CD46 (for *CD46*KO line). As shown in Fig. 3A-C, there is increased C3c *and* C4d in CM-HUS compared to healthy controls across all three cell lines. Deficiency of CD46 but not PIGA increases C4d deposition as compared to WT cells (Fig. 3A). The significant change in bioluminescence on the *CD46*KO line in samples from CM-HUS, as compared to the relative absence of change in healthy control samples, supports differential regulation of the CP in CM-HUS and identifies specificity of the *CD46*KO line for this disease. The critical role for CP vs AP control was further confirmed by examining the ability of sutimlimab or FDi (ACH-5548) to control either C3 or C4 deposition. Across all cell lines, both C3 and C4 are more consistently blocked by sutimlimab than FDi, suggesting a critical role for the CP (Fig. 3 D-L). These findings were also confirmed on WT TF-1 cells, the cell line used for the traditional mHam (Fig. S4). Furthermore, the changes in complement deposition pattern (increased C4d and C3c) are consistent between WT cells and the knockout cell lines lacking complement regulators. In summary, these results suggest CD46 is more important than CD55 in regulation of the CP in CM-HUS, and this assay identifies a physiologic stimulus inherent to the disease as opposed to creating an artifactual finding.

**Fig. 3.**
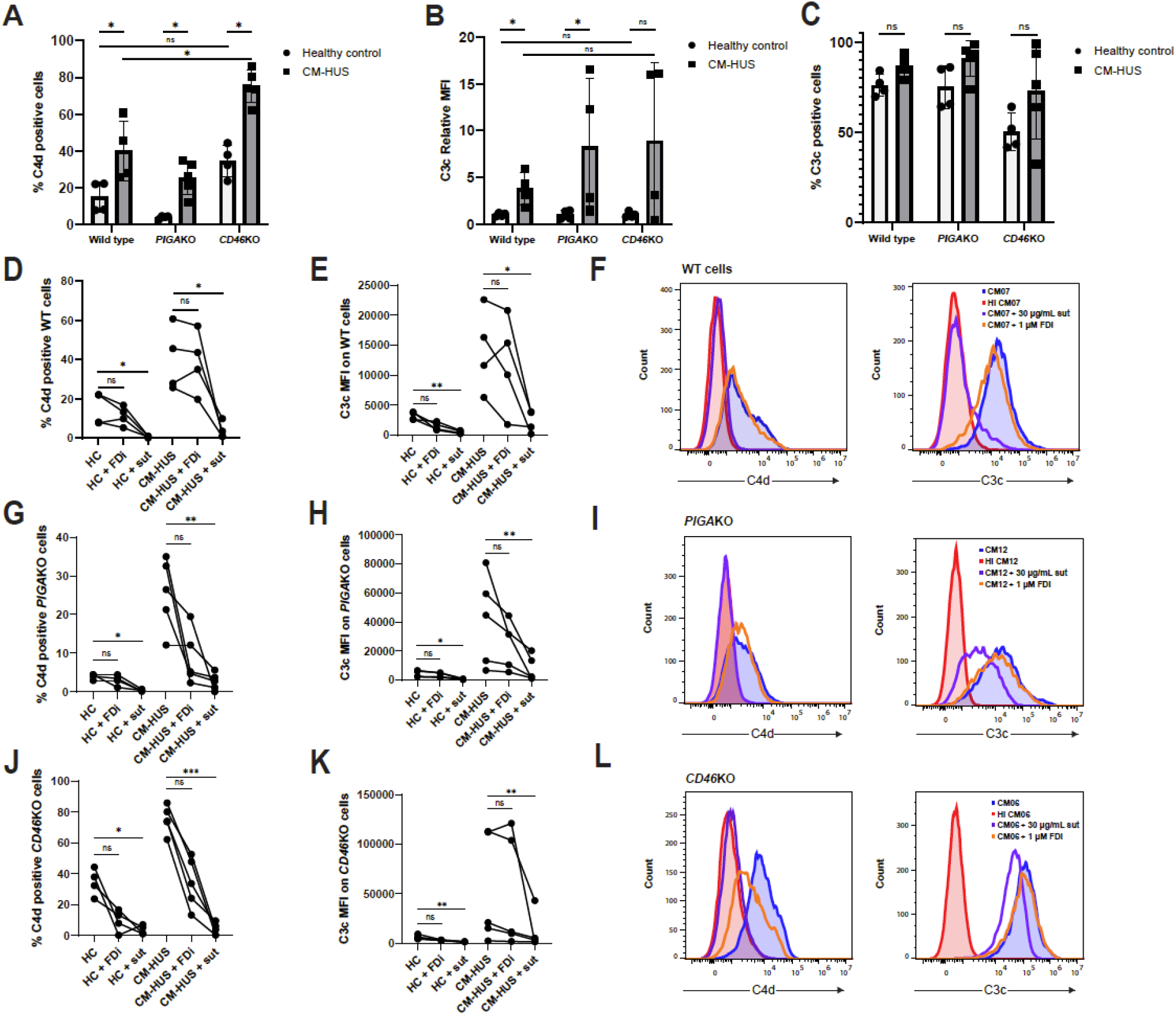
C4d and C3c deposition and inhibition in HC and CM-HUS samples. **A-L**, All experiments were done in the presence of C5 inhibitor, eculizumab (100 μg/mL). WT, *PIGA*KO, or *CD46*KO cells were treated with 20% serum in GVB^++^ for 30 minutes and then stained singly for either C4d or C3c and deposition quantified by flow cytometry. The same healthy control (n=4) and CM-HUS (n=4 for WT, n=5 for *PIGA*KO and *CD46*KO) were used across all cell types. ns, not significant, *p < 0.05, **p < 0.01, ***p < 0.001, ****p < 0.0001. **A**, Comparison of C4d deposition across cell types by percent positive cells determined by gating on heat-inactivated HC sample. **B**, Relative median fluorescence intensity (MFI) of C3c deposition on sample compared to average MFI of 4 healthy controls samples. **C**, Comparison of C3c deposition across cell types by percent positive cells determined by gating on heat-inactivated HC sample. **A,B,C**. Intra cell line differences between HC and CM-HUS *P* values were calculated with unpaired Mann-Whitney test. Inter cell line *P* values were calculated with a paired, two-tailed student’s t-test. **D,G,J**, C4d deposition on WT cells (**D**), *PIGA*KO (**G**), *PIGA*KO (**J**) in HC or CM-HUS with or without addition of sutimlimab (30 μg/mL, sut) or ACH-5548 (1 μM, FDi). **E,H,K**, C3c median fluorescence intensity (MFI) on WT cells (**D**), *PIGA*KO (**G**), *PIGA*KO (**J**) in HC or CM-HUS with or without addition of sutimlimab (30 μg/mL, sut) or ACH-5548 (1 μM, FDi). **D,G,J,E,H,J,K**, *P* values calculated using Friedman’s test for Dunn’s multiple comparisons. **F**, Example histograms of CM07 on WT cells with or without heat inactivation (HI), sutimlimab or factor D inhibitor. **I**, Example histograms of CM12 on *PIGA*KO cells with or without heat inactivation (HI), sutimlimab or factor D inhibitor. **L**, Example histograms of CM06 on *CD46*KO cells with or without heat inactivation (HI), sutimlimab or factor D inhibitor.

### *PIGA*KO and DKO biosensors can be used to identify patients who may benefit from increased dosing of C5 inhibitors or C1-INH

There are no standardized assays to evaluate membrane surface protection with the use of complement inhibitors^40–43^.The ability of the *PIGA*KO and DKO lines to identify a range of “normal” complement activity means these biosensors are potentially useful in monitoring therapeutic complement blockade. As shown in Fig. 4A,B, when on therapeutic complement inhibition, patient serum yields similar cell killing to that of heat inactivated samples on either the *PIGA*KO or DKO lines. Evaluating 14 samples from patients on eculizumab or ravulizumab showed an average 1 h relative luminescence of 91.7% (95th% CI 82.9-100.4) with only one patient with relative luminescence less than 88% (CM21 at 40.3%) (Fig 4C). Addition of either sutimlimab or eculizumab (but not FDi) to CM21 sera shows that additional cell protection could be offered by increasing complement inhibition (Fig. 4D). We also confirmed a dose dependent response for classical pathway inhibitors for sutimlimab, eculizumab, and C1-inhibitor (Fig 4E-G). Finally, we show that patient sera can be diluted with NHS to give a “titer” required to overcome the complement inhibition and show an activating stimulus even when on complement inhibitor therapy (Fig. 4H).

**Fig. 4.**
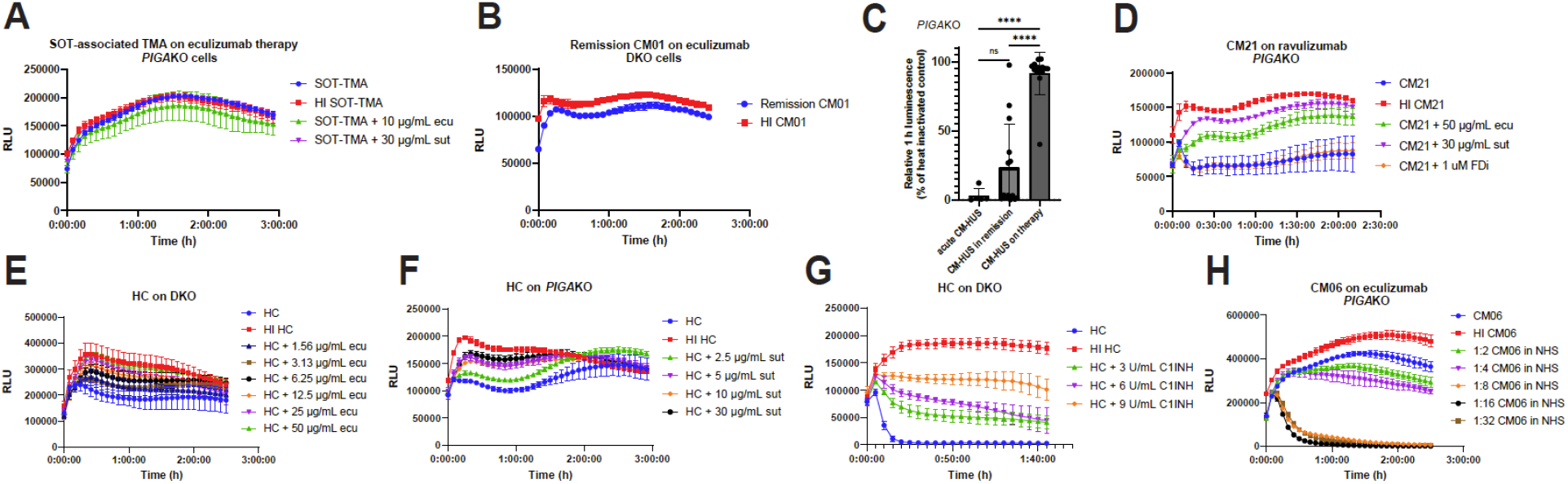
*PIGA*KO and DKO can be used to monitor therapeutic complement inhibition. **A**, Example tracing of solid organ transplant-associated TMA sample on therapeutic eculizumab ran on *PIGA*KO cells with or without addition of sutimlimab or eculizumab. **B**, Example tracing of CM01 remission sample after being started on eculizumab on DKO cells. **C**, Summary of relative luminescence at t = 1 h for CM-HUS samples (n=14). Unpaired acute and remission samples are included as a reference. *P* values were calculated using one-way ANOVA for Dunnett’s multiple-comparisons test. ns, not significant, *p < 0.05, **p < 0.01, ***p < 0.001, ****p < 0.0001. **D**, Example tracing of CM21 on ravulizumab therapy with or without the addition of additional C5 inhibitor (eculizumab), sutimlimab, or factor D inhibitor on *PIGA*KO cells. **E**, Example tracing of HC on DKO with increasing concentrations of eculizumab. **F**, Example tracing of HC on PIGAKO with increasing concentrations of sutimlimab. **G**, Example tracing of HC on DKO with increasing concentrations of C1-inhibitor. **H**, Acute CM-HUS sample on eculizumab therapy, CM06, on *PIGA*KO diluted with pooled normal human serum. **A,B,D,E,F,G,H**, Example traces plotted as mean +/- SD for each triplicate. RLU = relative luminescence units.

### IgM is the immunoglobulin species contributing to CM-HUS complement activation

Finding a predominant role for CP, we hypothesized this stimulation would be mediated by immunoglobulin. We utilized cleavage of IgG with IdeS (Fig. S5) or selective reduction of IgM with 3 mM dithiothreitol (DTT) to investigate the contributions of IgG or IgM, respectively. IdeS is an IgG-specific endopeptidase which cleaves human IgG into F(ab′)2 and Fc fragments; although F(ab’)2 is still available to bind to antigen, the loss of the complement-fixing Fc domain prohibits IgG-mediated complement activity^44^. DTT treatment is commonly used to differentiate IgG from IgM-mediated pathologic effects^45–47^. We confirmed DTT treatment causes a selective reduction of IgM with minimal effect on IgG using a non-reduced, unheated SDS-PAGE gel (Fig. 5A) and showed dose-dependent effects on complement inhibition in healthy control sera (Fig. S6). Given DTT has nonspecific effects on complement activation^48^, we also performed fractionated antisera (principally IgG) sensitization of sheep erythrocytes followed by quantification of CH50 with or without the presence of DTT. Under these conditions, there was an average CH50 reduction of ~35% at either pretreatment (3 mM DTT) or final concentrations of DTT (0.6 mM DTT) utilized in the bioluminescent mHam (Table S1). Additionally, demonstrating the strength of the assay in understanding complement activation pathways relevant to an individual’s disease presentation, we identified a patient with acute fatty liver of pregnancy and HELLP (Table S2) where activation is blocked with heat or eculizumab but not sutimlimab or DTT (Fig. 5B). Taken together these results demonstrate that DTT inhibitory effect in the assay is driven principally by reduction of IgM as opposed to nonspecific effects on serum or complement.

**Fig. 5.**
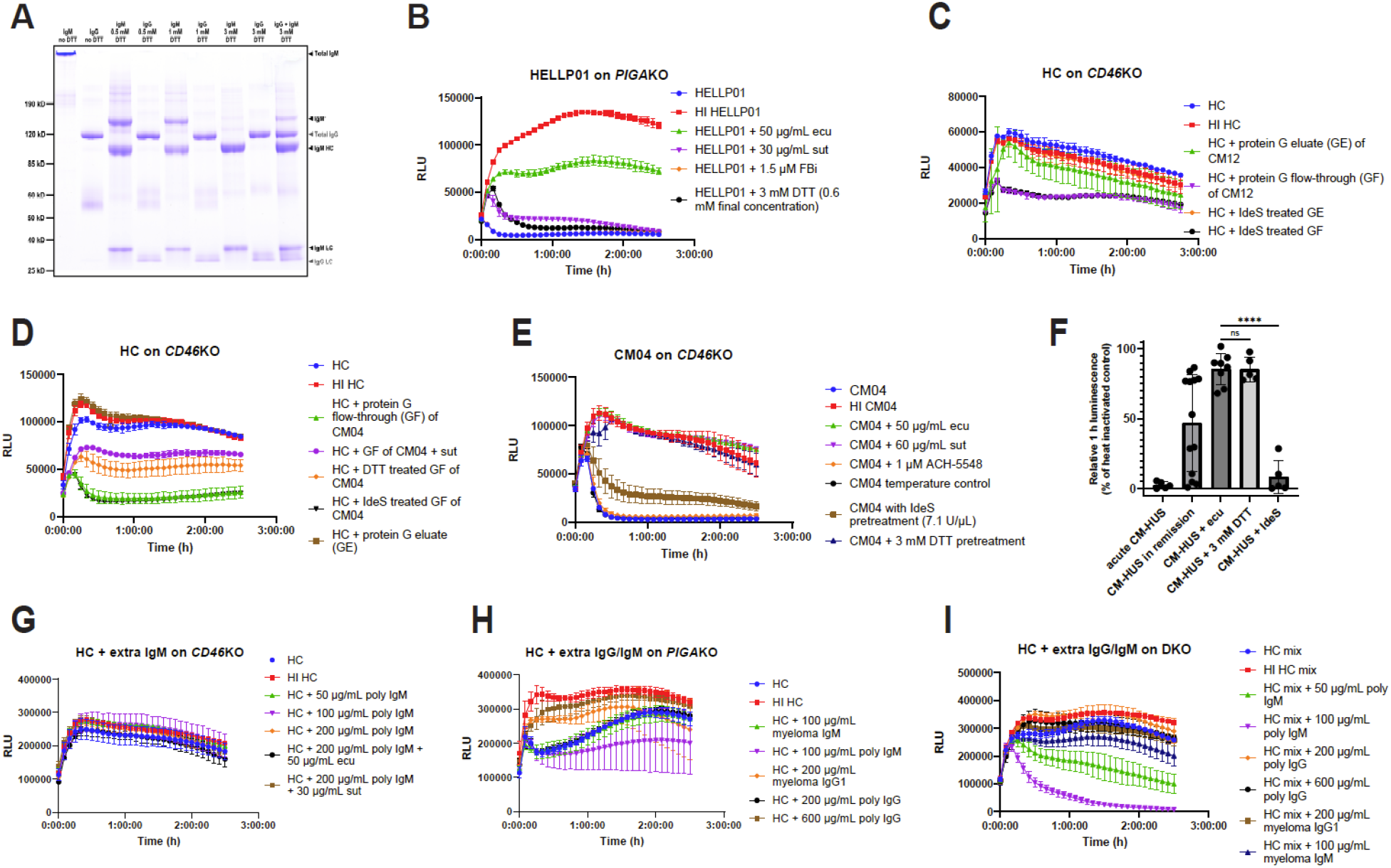
Serum treatment with DTT or IdeS and protein G spin columns identify IgM as the predominant immunoglobulin species leading to complement activation. **A**, Non-reducing, no heat SDS-PAGE gel of IgG and IgM with increasing concentrations of DTT. IgG (3 μg), IgM (3 μg), or both were incubated with DTT (0 mM, 0.5 mM, 1 mM, 3 mM) in PBS (pH 7.4) in a final reaction volume of 10 μL at room temperature for 30 min. Following treatment, samples were diluted with 4x loading dye (without reducing agent) and left at room temperature for 10 min prior to running SDS-PAGE gel and staining with coomassie. HC = heavy chain, LC = LC. IgM’ represents a partially reduced form of IgM. **B**, Sera from patient with hemolysis, elevated liver enzymes, low platelets (HELLP) with or without the addition of additional C5 inhibitor (eculizumab), sutimlimab, DTT, or factor D inhibitor on *PIGA*KO cells. **C**, CM12 sera (60 μL per column, 120-240 μL total) was processed over a protein G spin column to purify IgG (protein G eluate, GE) or isolate flow-through (protein G flow-through, GF). GE and GF underwent centrifugal concentration (as required), desalting, and buffer exchange into PBS. One third of final preparation per well was added to healthy control sera in the bioluminescent mHam on *CD46*KO cells with or without IdeS treatment (3.6 U/μL for 2 h at 37 °C). **D**, CM04 sera (240 μL) prepared as per **C**. Following preparation GF samples were treated with IdeS (3.6 U/μL for 2 h at 37 °C), DTT (3 mM pretreatment, final 0.6 mM), or sutimlimab (30 μg/mL). One third of final preparation (60 μL original volume) per well was added to healthy control sera in the bioluminescent mHam on *CD46*KO cells. **E**, Example tracing of CM04 in bioluminescent mHam on *CD46*KO cells with or without heat inactivation, temperature control (untreated serum heated to 37 °C for 30 min), eculizumab, sutimlimab, IdeS pretreatment, or DTT pre-treatment. Final concentrations listed except for DTT. **F**, Summary of relative luminescence at t = 1 h for CM-HUS samples treated with DTT (n=5) or IdeS (n=5). Unpaired acute, remission samples, and eculizumab spiked samples are included again as a reference. *P* values were calculated using one-way ANOVA for Dunnett’s multiple-comparisons test. ns, not significant, *p < 0.05, **p < 0.01, ***p < 0.001, ****p < 0.0001. **G**, Healthy control sera spiked with polyclonal IgM (extra 5, 10, or 20 μg/well) with final concentrations as indicated and ran in bioluminescent mHam on *CD46*KO cells. **H**, Healthy control sera spiked with myeloma IgM (10 μg/well), polyclonal IgM (10 μg/well), myeloma IgG1 (10 μg/well), polyclonal IgG (10 μg or 60 μg/well) with final concentrations as indicated and ran in bioluminescent mHam on *PIGA*KO cells. **I**, Healthy control sera spiked with polyclonal IgM (5 or 10 μg/well), polyclonal IgG (20 or 60 μg/well), myeloma IgG1 (20 μg/well), or myeloma IgM (10 μg/well) with final concentrations as indicated and ran in bioluminescent mHam on DKO cells. **B,C,D,E,G,H,I**, Example traces plotted as mean +/- SD for each triplicate. RLU = relative luminescence units.

Isolation of CM-HUS IgG with protein G spin columns followed by addition to healthy control serum consistently showed increased activity in the flow-through as opposed to the IgG eluate fraction (Fig. 5C). Treatment of flow-through with DTT or sutimlimab partially blocks these effects whereas IdeS treatment does not (Fig. 5D). Moreover, pretreatment of CM-HUS sera with DTT but not IdeS results in almost complete blockade of observed activity (Fig. 5E,F) with a mean 1 h relative luminescence of 85.4% (mean 95% CI 74.5-96.3%) vs 8.3% (mean 95% CI 6.7%-23.3%) for DTT and IdeS, respectively.

We next investigated the ability of commercial preparations of polyclonal IgG, polyclonal IgM, monoclonal IgG, and monoclonal IgM to activate complement when spiked into healthy control sera across all three cell lines. For the DKO line, a pool of healthy control sera was selected for low basal activity. As seen in Fig. 5G, polyclonal IgM cannot stimulate activation on the *CD46*KO line. There is a trend towards activating on the *PIGA*KO line (Fig. 5H), but the effect is clearest on the DKO line (Fig. 5I), where there is a dose dependent response with IgM but not monoclonal IgG/IgM or polyclonal IgG. This activation is again blocked by both eculizumab and sutimlimab (Fig. S7).

Innate, polyreactive IgM demonstrates low affinity interactions for a wide variety of substrates^49,50^. The ability of polyclonal IgM and healthy control sera to activate complement on a physiologic membrane surface provides evidence that this is the culprit immunoglobulin species. However, there is a clear difference in relative affinity of the polyreactive IgM for the cell surface even among healthy controls as evidenced by a range of activity on the *PIGA*KO line from 0% to 100% and that some healthy controls show only minimal complement activity even on the DKO cell line–but can be stimulated with the addition of polyclonal IgM preparations. Furthermore, unlike the activity observed in CM-HUS, neither healthy controls nor polyclonal IgM preparations have significant activity on the *CD46*KO line suggesting a difference in total complement fixing avidity. As a preliminary evaluation of this concept, we examined the degree of IgG and IgM binding in a selection of CM-HUS and healthy control samples. There was a trend towards increased IgG and IgM staining in CM-HUS compared to healthy controls (Fig. 6A). Using a relative MFI of >95th percentile of healthy controls (>2.05 for IgG and >1.36 for IgM) as a positive threshold, there was no difference in IgG staining (0 of 9 for HC vs 3 of 17 for CM-HUS, p = >0.53, Fisher’s exact test), but significantly more positive samples for IgM in CM-HUS compared to HC (0 of 7 for HC vs 8 of 15 for CM-HUS, p = 0.0225, Fisher’s exact test). Furthermore, in the CM-HUS samples increased IgM staining is more frequently seen in acute (4 of 6) as opposed to remission samples (4 of 9). Finally, IgM but not IgG staining can be reduced by treatment with DTT (Fig. 6B,C).

**Fig. 6.**
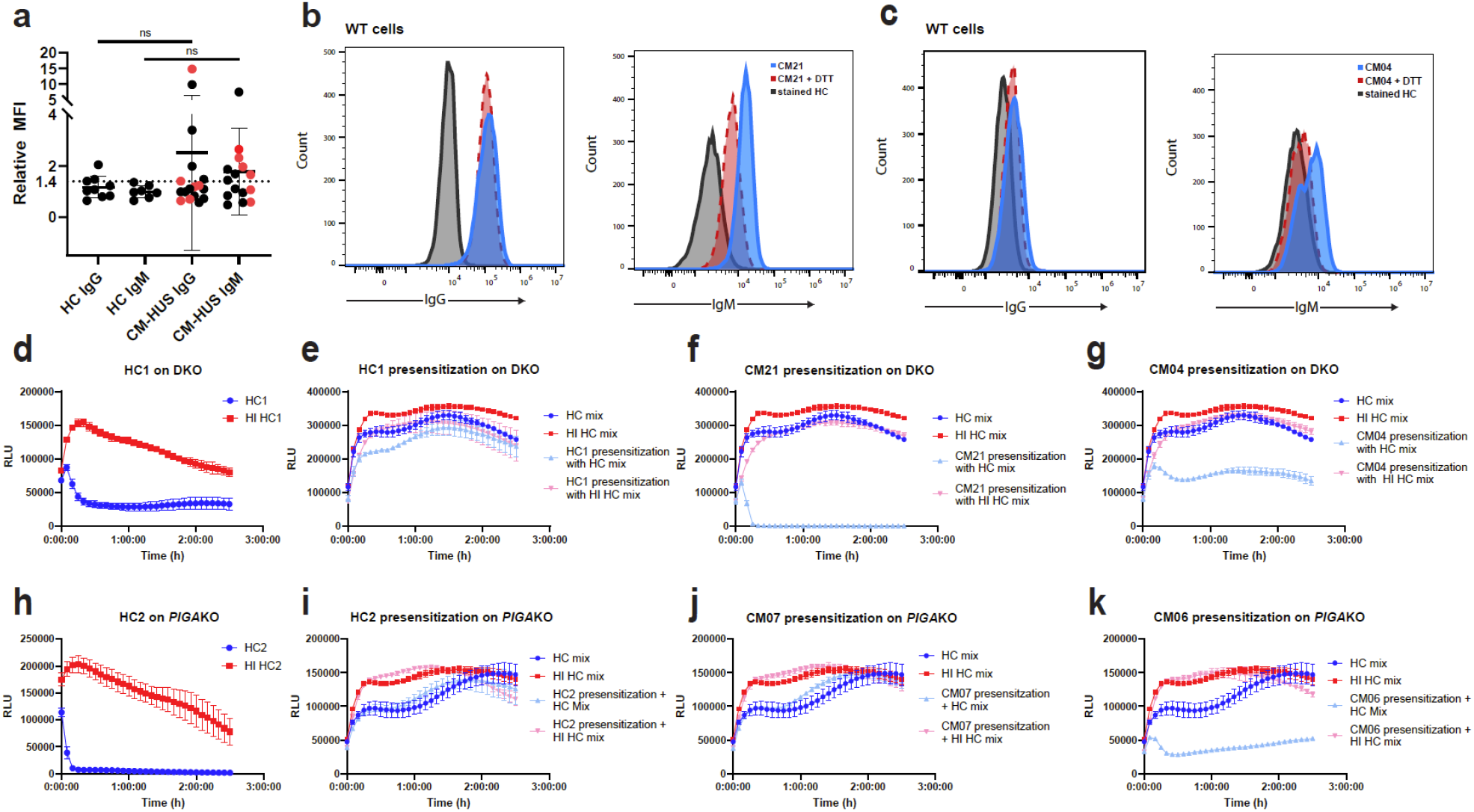
CM-HUS sera has increased IgM binding to HEK293 cell surfaces and can pre-sensitize cells even after heat inactivation of serum. Heat inactivated sera was utilized for all cytometry and presensitization experiments to eliminate preloading of cells with C3 or C4 fragments. **A**, WT cells were incubated for 45 min at 37 °C with either heat-inactivated HC or CM-HUS sera (20% in GVB_++_), washed, and evaluated for deposition of IgG (HC n=9, CM-HUS n=17) or IgM (HC n=7, CM-HUS n=15) by flow cytometry. Acute CM-HUS samples (n=6) indicated by red dots. MFI = median fluorescence intensity. Relative MFI calculated as ratio of sample MFI compared to average of 4 healthy controls ran with each experiment. *P* values were calculated using unpaired, two-tailed t test with Welch’s correction. ns, not significant. **B**, Example histograms of IgG (left) or IgM (right) staining from CM21 on WT cells with (red) or without (blue) pretreatment of serum with DTT (3 mM pretreatment). HC stained IgG or IgM stained cells included as baseline reference population (grey). **C**, Example histograms of IgG (left) or IgM (right) staining from CM04 on WT cells with (red) or without (blue) pretreatment of serum with DTT (3 mM pretreatment). HC stained IgG or IgM stained cells included as baseline reference population (grey). **E**-**G,I**-**K**. Specially selected low activity healthy control sera mix was utilized to facilitate complement activity in the bioluminescent mHam after either DKO (**E**-**G**) or *PIGA*KO (**I**-**K**) cells were pre-sensitized (20% sera treatment for 45 min in DMEM), washed, and resuspended in GVB_++_ then ran in the bioluminescent mHam. Baseline activity of the healthy control mix sera shown in each tracing as dark blue (HC sera mix) and dark red (HI HC sera mix) **D**, Bioluminescent mHam tracing of HC1 on DKO. **E**, Bioluminescent mHam tracing of HC1 presensitized DKO cells with HC sera mix. **F**, Bioluminescent mHam tracing of CM21 presensitized DKO cells with HC sera mix. **G**, Bioluminescent mHam tracing of CM04 presensitized DKO cells with HC sera mix. **H**, Bioluminescent mHam tracing of HC2 on *PIGA*KO. **I**, Bioluminescent mHam tracing of HC2 presensitized *PIGA*KO cells with HC sera mix. **J**, Bioluminescent mHam tracing of CM07 presensitized *PIGA*KO cells with HC sera mix. **K**, Bioluminescent mHam tracing of CM06 presensitized *PIGA*KO cells with HC sera mix. **E**-**K**, Example traces plotted as mean +/- SD for each triplicate. RLU = relative luminescence units.

### Pre-sensitization of PIGA or DKO cells with CM-HUS sera and then assessing ability to cause activation in healthy control sera demonstrates cell-bound IgM alone is sufficient to stimulate complement activation

As a final method to explore the differential complement-fixing avidity of CM-HUS sera, we pre-sensitized either DKO or *PIGA*KO cells with the heat-inactivated CM-HUS or healthy control sera. The cells were washed and then incubated with healthy control sera (as a source of complement). As shown in figure 6 (and Fig. S8), pre-sensitization with acute CM-HUS serum (n=2) significantly increased complement activity of the healthy control sera whereas only 1 of 7 CM-HUS remission samples similarly tested was positive. The positive remission sample was CM21 which also had incomplete blockade on ravulizumab. Along with the trend of increased IgM binding in acute compared to remission CM-HUS samples (*vide supra*, Fig. 5), these results suggest a shift in the IgM pool in the acute setting of CM-HUS that resolves once in remission. Importantly, given heat inactivation, washing, and use of healthy control serum in the assay, this also demonstrated that IgM alone was sufficient to stimulate pathologic complement activity in some cases of CM-HUS. Together these results suggest a role for both “natural, polyreactive” and “autoreactive” IgM in the disease^51^.

## Discussion

We have developed complement biosensors capable of detecting surface-directed complement activation in real time. In contrast to fluid phase markers of complement (sC5b-9, Bb or Ba), cell-directed complement activation is more specific to CM-HUS disease pathogenesis^10,11,31^. These biosensors are unique among complement diagnostics in that they allow for detection of a range of complement activity without antibody sensitization^52,53^ or cell stimulation^11^ allowing for easy discrimination of activity coming from patient serum as opposed to the sensitization technique.

Since 1974 depression of C3 more than C4 in CM-HUS has been noted, and even at that time in accordance with our own observations it was shown that the “C3-splitting activity” was greater in all pathway buffer and suggested a classical pathway stimulus in the disease^54^. A CP stimulus from a polyreactive IgM profile provides a mechanistic explanation for both disease activity, even in those without variants in complement regulatory proteins, and the variable penetrance of the disorder in carriers of known pathogenic variants. It also explains the common propensity for C1q, C4d, and IgM deposition in histologic specimens of thrombotic microangiopathy, including deltoid skin biopsies^55–64^. These tissue findings contrast with what is seen in aHUS plasma where C3 is depressed in ~22-32% of samples compared to only 6.7% for C4^65^. The reason for a higher incidence of low C3 is likely explained by the role for C4 as a complement initiating protein versus C3 as an amplifying protein; this is confirmed in quantitative radioisotopic studies which show that after CP stimulus, 30x more molecules of C3 than C4 are deposited on the cell surface^66^. Prior studies have shown that 80% of complement activity comes from alternative amplification on a solid surface^67,68^. Our assay data showing ongoing CP activity even in a patient with C3 depletion as well as activity with triple AP blockade provides further evidence that in the face of overwhelming CP stimulus, the role for AP amplification is less significant ^29,30,69^.

Atypical HUS has previously been implicated as an autoimmune disease with a role for autoreactive antibodies^70–72^. Leung et al implicate anti-endothelial cell antibodies in causing complement-mediated cytolysis in the disorder^70^. More importantly, their data provide additional evidence that the CP stimulus is relevant to endothelial surfaces in CM-HUS. In contrast to our data, they implicate both IgG and IgM in endothelial injury, which cannot be entirely excluded by our studies on HEK293 cells that have a different repertoire of cell-surface antigens. Similar findings in the traditional mHam/TF-1 cell line again suggest the generalizability of these findings across cell surfaces and reinforce that the stimulus is not directed at a specific cell epitope, but rather driven by low affinity, polyreactive interactions.

Our study highlights many important observations on the role of complement and IgM in disease pathophysiology. Although most literature focuses on their role as AP regulators, there is also evidence that FH and CD46 mitigate classical pathway activity^34–38,73–75^. Our data suggest that in CM-HUS the role of CD46 in regulating classical activity is more important than its role in controlling the amplification loop. Interestingly, in C3 glomerulopathy, another disorder characterized by alternative pathway variants, the disease can rarely present or even transition from dense deposit disease with predominantly C3-containing deposits to immune complex deposits^76–78^. There is also well-known interplay between polyreactive IgM and FH. In mice, FH is suspected of functioning more akin to CR1 in primates and has been shown to be crucial to controlling immune complexes injury in the glomerulus^79,80^. Furthermore, FH deficiency in mice exacerbates the ability of polyclonal and monoclonal IgM recognizing phospholipids to deposit on kidney mesangial cells, leading to complement activation and cell damage^81^. In human studies, low C4 and elevated C4d in an aHUS cohort with FHAA correlated with anemia and IgM on kidney biopsy^59^. In lupus nephritis, IgM independently contributes to C3 deposition, and those with the highest IgM staining and lowest FH levels show exacerbated proteinuria and depressed systemic C3 levels^82^. Additionally, IgM as opposed to IgG has been shown to be a more important complement stimulus in xenotransplantation^83,84^. Lastly, in COVID-19, a disorder with known abnormal complement activation, polyclonal IgM has been implicated in disease severity^85^.

Our study has several limitations including underrepresentation of pathogenic variants (~25-38% in our study compared to 40-60% in larger cohorts) and low representation of those with factor H variants in particular. However, compared to those with a detectable variant, the CP stimulus is more relevant to explaining disease pathology in those *without* a known AP variant, which comprises up to half of all CM-HUS. Larger studies are needed for validation and to correlate ongoing positivity in the assay with relapse and non-relapse related outcomes. As an added caveat, since fluid phase and membrane-directed complement activation likely lie on a spectrum, the diagnostic accuracy of the test could be decreased in cases of strong fluid-phase activation leading to complement depletion.

In summary, our data with biosensors reveals that CM-HUS is a multi-hit disease with a CP stimulus, addressing the mystery of why 40% of CM-HUS lack complement specific variants or autoantibodies. We demonstrate this finding even in patients with pathogenic variants in cell surface complement regulators such as membrane cofactor protein (MCP, CD46). Variants in complement regulatory genes (FH, CD46, FB, C3, FI) lower the threshold for disease but are not necessary or sufficient to cause disease. CP activation in the disease is at least partially driven by polyreactive or autoreactive IgM and is readily inhibited by DTT or inhibition at C1s or C5. Lastly, we show that the bioluminescent mHam provides a rapid assay to help distinguish CM-HUS from TTP and can be used to monitor complement inhibition. Taken together, these data suggest a breakdown in IgM immunologic tolerance as a key driver of CM-HUS.

## Acknowledgements

We thank S. Peter Howard (University of Saskatchewan, Saskatoon, Canda) for gracious donation of the pro-aerolysin utilized in this study. We thank Alexion Pharmaceuticals (Boston, MA) for generously providing both factor D inhibitors, ACH-4471 and ACH-5548. M.A.C. was supported by the J. Mario Molina Physician-Scientist Scholarship and NIH T32 Training Program in Hematology (T32HL007525). This work was funded by N.H.L.B.I (R56HL133113) and the Department of Defense (W81XWH2110898). The funders had no role in study design, data collection and analysis, decision to publish or preparation of the manuscript.

