## Supplemental Methods and Figures for "Complement Biosensors Identify a Classical Pathway Stimulus in Complement-Mediated Hemolytic Uremic Syndrome"

Inclusion criteria and sample collection
This study was performed leveraging existing CM-HUS samples. SST/gold top samples were previously processed and stored at -80°C as part of the Hopkins Complement Associated Disorders Registry with samples collected from 2014-2023 (NA_00090162). Samples were processed and frozen within 3 hours of collection and stored in multiple aliquots to avoid repeated freeze/thaws. All patients underwent written/informed consent. All samples from the registry were included to be ran on the bioluminescent mHam if they 1) have a clinical diagnosis of CM-HUS or aHUS by an inpatient or outpatient consulting hematologist, 2) had not yet received plasma exchange or eculizumab therapy, 3) sufficient sample remained in the biorepository to run the sample. Given an abundance of samples for patients already on eculizumab or ravulizumab in the registry, these samples were selected at random for inclusion. TTP was diagnosed by an ADAMTS13 activity level less than 10%. STEC-HUS was excluded in patients who presented with diarrhea. Samples were defined as acute if they were collected within 14 days of initial manifestations, 2) had not yet received either plasmapheresis/plasma exchange, C5 inhibitory therapy or caplacizumab, and 3) had not yet achieved hematologic remission (defined as Hgb >11 g/dL or recovery to baseline, LDH within normal limits and platelet count >150x$10^3$/µL). Samples from patients with renal-limited TMA (ie without systemic or hematologic manifestations) were excluded. Healthy controls were non-hospitalized, non-pregnant, healthy volunteers aged 18-65.

Description of Key Reagents
Livelight HEK293 cell line was from 490 BioTech (Knoxville, TN). Proaerolysin was obtained from Peter Howard (University of Ottawa, Saskatoon, Candada). Sutimlimab and eculizumab were obtained from hospital pharmaceutical waste. Factor D inhibitors ACH-5548 and ACH-4471 were obtained from Alexion Pharmaceuticals (Boston, MA). Factor B inhibitor iptacopan and C3 inhibitor compstatin were from MedChemExpress (Monmouth Junction, NJ). All inhibitors were tested for activity in the Wieslab AP or CP activity kits from Eagle Biosciences/Svar (Amherst, NH). Iptacopan, ACH-4471, ACH-5548 were dissolved in DMSO (10 mM for iptacopan and ACH-4471 or 20 mM for ACH-5548) for storage. Any subsequent dilutions for all inhibitors were in PBS. Longterm storage was at –80 °C and short-term storage of working aliquots was at –20 °C. Unsensitized sheep erythrocytes, $GVB^{++}$, $GVB^0$, VBS, MgEGTA were from Complement Technologies (Tyler, TX). Pooled NHS were from both Complement Technologies (Tyler, TX) and QuidelOrtho (Athens, OH). Heat aggregated gamma globulin was from QuidelOrtho (A114). Antibodies included: monoclonal mouse anti-human C4d (QuidelOrtho, cat #A213), rabbit polyclonal FITC-conjugated C3c (abcam, cat# ab4212), AF647- conjugated goat anti-Human IgG (H+L) (Thermo fisher, cat# A21445), AF488-conjugated goat anti-Human IgM (Heavy chain) (Thermo fisher, cat#A21215), PE-conjugated mouse anti-human CD46 (molecular probes, cat#A-15776, clone MEM-258), and BV421-conjugated mouse anti-human CD55 (BD optibuild, clone IA10), and BV510-conjugated mouse anti-human CD59 (BD optibuild, clone H19). Secondary for C4d was AF488-conjugated goat anti-Mouse IgG (H+L) (cat# A32723). Polyclonal IgG (cat#AG711), myeloma IgG1 (cat# 400120), and myeloma IgM (cat#401108) were from EMD Millipore, and native polyclonal IgM was from BioRad (cat# 5275-5504). IdeS protease was from Promega (cat#V7511). Anti-sheep red blood cell stroma antibody produced in rabbit was from Sigma-Aldrich (cat#S1389). Purified cas9 protein and sgRNA sequences were from Synthego (Redwood City, CA). CD46 sgRNA sequences were CD46 target#1: CAATTGTGTCGCTGCCATCGAGG and CD46 target#: CGATTTCCAGTAGTCGAAAATGGG. Check primers were left- ATCTTGCATTCCATTCCTTGTC and right- TTAAGACACTTTGGAACTGGG. All designed with CHOPCHOP[1].

AP Buffer
Alternative pathway buffer (GVB-MgEGTA) was prepared fresh for each assay by diluting 650 uL MgEGTA (0.1 M $MgCl_2$, 0.1 M EGTA, pH 7.3) to 5 mL with $GVB^0$ (both from Complement Technologies). Final concentration 13 mM Mg and EGTA. pH was adjusted to 7.30-7.45 with dropwise addition of HCl (6 M).

Cell culture and propagation
Each cell line was cultured and maintained in DMEM with 9% FBS, 90 µg/mL penicillin/streptomycin, 1x glutaMAX). Cells were maintained in log phase growth by passage every 2-3 days. Cell lines were

replaced with an early passage line approximately every 3 months. Cells were either passed the day prior to the assay (preferred) or rarely cells were utilized for the assay after passage two days prior running the assay if cell density was still <80% confluent with good cell morphology and viability after washing was >92% (as a minimum requirement for utilizing the cells in the bioluminescent mHam). Wash buffer was Hank's Balanced Salt Solution without calcium or magnesium unless otherwise indicated. Cell lines were tested periodically to be mycoplasma free using PCR based Venor GEM Mycoplasma detection kit (Sigma-Aldrich, MO). Viability was assessed by trypan blue staining using a Denovix celldrop cytometer.

Cell line Engineering

A *PIGA*KO HEK293 clone was selected by treatment of ~10 million Livelight HEK293 cells with 0.5 nM proaerolysin in DMEM (9% FBS, 1x penicillin/streptomycin, 1x glutaMAX). Media was exchanged following the first 24 hours of treatment to remove nonadherent cytolytic cell fragments and cells were allowed to propagate for an additional 2-3 weeks with ongoing proaerolysin selection. Following 2-3 weeks of treatment individual adherent colonies were carefully removed and propagated separately. CD55 and CD59 negativity were confirmed by flow cytometry and a final treatment with 1 nM proaerolysin. A CD46 knockout line (*CD46*KO, LL HEK293 $PIGA^{wt}$/$CD46^{-/-}$), or double knockout (DKO, LL $PIGA^{-}$/$CD46^{-/-}$) were created by nucleofection of preformed RNP with two separate sgRNA sequences on the WT cell line or *PIGA*KO cell line. Briefly for each cell line, Cas9-RNP complex was preformed by mixing 3 µL sgRNA target#1 (100 pmol/µL in TRIS-EDTA), 3 µL sgRNA target#2 (100 pmol/µL in TRIS-EDTA), and 5 µL spCas9 (20 µM from synthego) in Lonza nucleofection solution plus supplement (6 µL) and incubated on ice for 10-15 minutes. Cells (800k per condition) were harvested with trypsin/EDTA, neutralized with DMEM with 9% FBS, washed in HBSS with careful removal of excess buffer, and then resuspended in prewarmed nucleofection buffer plus supplement (100 µL). Cells were added to preformed RNP complex, transferred to a 100 µL nucleovette, and nucleofection performed on a Lonza 4D (program CM-130). Immediately following nucleofection cells were transferred to a pre-warmed media in a 25 cc tissue culture flask. Following approximately one week of propagation, the mixed polyclonal population was single cell sorted by selection on the CD46 negative population (CD46-PE Ab at 1:25 dilution) at the JHH Bloomberg School of Public Health cell sorting facility. Following propagation, CD46 negativity was confirmed by flow cytometry and PCR with check primers to demonstrate expected fragment length differences.

Bioluminescent mHam

Cells were harvested after rinsing with PBS and brief treatment with trypsin 0.25%/EDTA 2.21 mM (~3 min treatment at room temp until detached). Trypsin was neutralized with FBS containing media and cells diluted in HBSS (10 mL). Cells were pelleted by centrifugation (1000 rpm for 7 min) and resuspended in culture media and kept at 37 °C until ready for use in the assay (typically used within 1-3 hours). Following cell harvest, serum samples were prepared by rapid thaw at 37 °C until only a trace quantity of ice remained and then allowed to finish thawing on ice. Aliquots of 66 uL (for untreated or inhibitor treated samples) or 70 uL (for heat inactivation) were created. Heat inactivated samples were treated in a 56 °C water bath for 30 min and then placed on ice. Inhibitors were diluted in PBS to 100x final assay concentration and added to serum aliquots at 5x final reaction concentration, eg, 3.47 µL of 100 µM ACH-4471 was added to 66 µL of serum, and a 20 µL aliquot of the inhibitor treated sample is added to 40,000 cells in $GVB^{++}$ (80 µL) as the final step in assay preparation. Final inhibitor concentrations (unless otherwise indicated in figure legends) were eculizumab (50 µg/mL, ~338 nM), sutimlimab (30 µg/mL, ~204 nM), ACH-4471 (1 µM), ACH-5548 (1 µM), iptacopan (1.5 µM), compstatin (18 µM), DTT (0.6 mM). Pretreatment of serum with DTT (3 mM) occurred at room temperature for 30 minutes and then sample was again placed on ice to slow ongoing reduction. Following serum preparation, the previously harvested cells were washed with HBSS, pelleted by centrifugation, and resuspended in prewarmed $GVB^{++}$ at a concentration of 2000-3000 cells/µL with final recount on a DeNovix celldrop cytometer. Final assay plate (96-well, white opaque flat-bottom) is prepared in triplicate for each condition by addition to each well by multichannel pipette of 1) 40,000 cells, 2) appropriate amount of prewarmed $GVB^{++}$ to bring volume up to 80 µL, and finally 3) addition of serum (20 µL) that has been allowed to sit at room temperature for 10-15 minutes. Heat aggregated IgG, Polyclonal IgG, polyclonal IgM, or myeloma IgG1 or myeloma IgM were used as supplied or reconstitution as per manufacturer's instructions. When immunoglobulin was added to the assay, it was added to healthy control sera known to have "low activity" on that cell line. The volume of GVB++ was reduced as appropriate (typical quantities ranged 5-60 µg with final concentrations shown in the figure legend). Similarly, heat aggregated IgG and immunoglobulins were added to the cells as opposed to the serum to avoid complement depletion prior to initiation of assay. After preparing the microtiter plate, luminescence was monitored by serial luminescence measurements (every 5 min at 37 °C) on a BMG ClarioStar luminometer. The 1 h relative luminescence was used as a standardized timepoint to compare complement activity. Relative luminescence was calculated as the ratio of the average of serum or inhibitor treated-serum to that of heat-inactivated control. "Positive" assay is defined as the lower 85th percentile on both lines: 12% for the *PIGA*KO and 62% for the *CD46*KO line. Descriptive and comparative statistics were performed in GraphPad Prism. *P* values were calculated using one-way ANOVA for Dunnett's multiple-comparisons test.

IdeS treatment of serum or IgG fractions

IdeS was obtained from Promega and reconstituted per manufacturer's instructions. Healthy control sera was used to optimize IdeS cleavage at either 3.6 or 7.1 U/µL at 37 °C. At predefined timepoints (0, 15, 30, 45, 60, 120, 180 min), 0.5 uL of this solution was diluted with 24.5 uL of ice-cold PBS. Samples were kept on ice until immediately prior to running SDS-PAGE gel at which point the sample (10.4 µL) was treated 4x non-reducing loading buffer (3.5 µL) and heated to 70 °C for 10 min prior to loading of entire sample on NuPage 4-12% Bis-Tris precast gradient gel and ran in NuPage MOPS running buffer followed by Coomassie staining. Following optimization, typical IdeS cleavage conditions were 7.1 U/µL (10 µL of 50 U/µL IdeS added to 60 µL serum) for 30 min at 37 °C, or if cleaving IgG alone typical conditions were 3.6 U/µL (final concentration IdeS) for 60-90 min at 37 °C. An untreated sample of serum was similarly kept at 37 °C without the addition of protease as a temperature control, but in no case did incubation of serum for 30 min at 37 °C result activity decrement on the bioluminescent mHam.

Pre-sensitization of cells with bioluminescent mHam

A pool of 2-3 healthy controls with low activity on the DKO cell line was created to act as a source of complement for the assay and designated as "HC mix" and all patient samples were heat-inactivated prior to incubation with the *PIGA*KO or DKO cell lines to prevent complement deposition during the presensitization step. *PIGA*KO or DKO cells were harvested, washed, and resuspended in DMEM without FCS (360,000 cells in 80 µL DMEM). Heat-inactivated sera (30 min at 56 °C) from acute CM-HUS (n=2), CM-HUS in remission (n=7), or HC (n=3) was added and incubated for 45 min at 37 °C in a CO2 controlled environment. Cells are diluted to 250 uL in HBSS, centrifuged (7 min at 1000 rpm). Cells are then washed 1x with HBSS (250 uL) and resuspended in GVB++ (160 uL). After a final cell count, cells are run in the bioluminescent mHam as previously described with low activity "HC mix" serum. Curve tracings are compared to that of HC mix and included for reference on each figure tracing.

Protein G spin column IgG purification

Spin columns and centrifugal protein concentrators were used for rapid purification of immunoglobulin with the aim of preserving complement activity in both the eluate and flow through fractions. Protein G spin columns (0.2 mL size), Zebaspin desalting columns (0.5 mL, 7k MWCO), wash buffers, binding buffer, and elution buffers were obtained from Thermo scientific (Rockford, Il). Vivaspin 500 5KDa MWCO centrifugal protein concentrators were from Cytiva (Uppsala, Sweden); all used per manufacturer's instructions. CM12, CM04 or HC sera (20 µL for each expected well with no more than 60 µL per column, 120-240 µL total) was diluted to 500 µL in binding buffer and loaded onto a protein G spin column to purify IgG (protein G eluate, GE) or isolate flow-through (protein G flow-through, GF). GE and GF underwent centrifugal concentration (as required) with Cytiva centrifugal protein concentrators, and finally desalting/buffer exchanged into PBS with a Zebaspin desalting column. Protein concentration was approximated on a nanodrop spectrophotometer. Quality of preparation was evaluated by SDS-PAGE; given known residual IgG in the flow through, these fractions were also treated with IdeS (3.6 U/µL for 2 h at 37 °C). For CM04, GF samples were also pretreated with 3 mM DTT for 30 min at room temperature. GE or GF samples were added to HC sera by equal distribution of volume (as opposed to protein amount), such that 20 uL of sera input volume was added per well. Samples were kept at 4 °C and used immediately after preparation on the same day.

Sheep erythrocyte sensitization and CH50

A custom, in house CH50 assay was designed based upon the methods of Mayer[2] and Morgan[3]to allow for sensitization with fractionated antiserum to sheep RBC stroma containing principally IgG as opposed to IgM sensitizing reagents. The fractionated antiserum was resuspended as per manufacturer's instructions. Briefly, unsensitized sheep erythrocytes (E) were obtained from CompTech. They were washed twice with VBS immediately prior to use in the assay with all centrifugations occurring at 1000*g* for 5 min. The E concentration was adjusted so that final OD490 ranged 0.5-0.7 with water lysis of 50 µL of E with 200 µL water. Appropriate hemolysin concentration to obtain near 100% lysis was obtained by titration of hemolysin reagent: 50 µL E were incubated with 25 µL of serial diluted hemolysis (1:10, 1:20, 1:30, 1:40, 1:50, and 1:100) and 25 µL of a 1:25 dilution of serum (1:50 final) in VBS at 37 °C with intermittent agitation for 30 min. Following incubation, wells were diluted with 150 µL ice cold VBS and absorbance read at OD490; concentration of hemolysin selected for use was the next more concentrated dilution after the dilution at which a 100% lysis plateau was achieved (typically 1:30-1:40). To perform the assay, E were sensitized with hemolysin at the concentration determined above in VBS for 30 min at 37 °C with intermittent agitation. Sensitized erythrocytes were washed twice with VBS and resuspended in the original volume of VBS and used immediately in the assay. HC sera was incubated with or without 3 mM or 0.6 mM DTT for 30 minutes and then subsequently diluted 1:50 in VBS. In the wells of a 96 well microtiter plate, a further series of dilutions in VBS of each 1:50 diluted serum sample was made: 1:10; 1:5; 1:4; 1:3: 2:5; 1:2; 5:2; 3:1; 4:1; 5:1; 10:1; 10:0; 50 uL/well. EA (50 µL) were added to each well. Control wells for each assay in duplicate included 100% lysis and cell blank in VBS. In addition to pretreated serum, DTT was also added in replace of VBS to achieve a final assay concentration of either 0.6 or 3 mM DTT. The plate was incubated at 37 °C for 30 min with intermittent agitation, control well lysed with addition of water to final volume of 250 uL. Ice-cold VBS to a final volume of 250 µL was added to the remainder of the wells; the plate was centrifuged and 200 µL supernatant placed in a new 96 well microtiter plate for final OD read at 515 nM. Percent hemolysis for each well was calculated based upon blank corrected data. A linear graph of the log of diluted serum volume in uL (x) vs log of [(y/(1-y)] was created and solved for log(x) when y=0, converted from logarithmic base to find K (CH50 unit), corrected for original 1:50 dilution (multiply by 50), and then converted to CH50 units per mL. The experiment was repeated on two separate days to ensure consistency and averaged. These CH50 units are distinct from those typically reported in standardized assays.

Flow Cytometry

C3c and C4d

All cells were harvested and washed as described for the bioluminescent mHam. 300-360k WT LL HEK293, *PIGA*KO, *CD46*KO, WT TF1 cells in $GVB^{++}$ (80 µL) were treated with eculizumab-spiked, healthy control or CM-HUS serum (20 µL, final eculizumab 100 µg/mL in 100 µL total volume) for 30 min at 37 °C. Additional conditions included heat inactivation (56 °C water bath for 30 min), sutimlimab (30 µg/mL), and ACH-5548 (1 µM, FDi). Cells were washed with FACS staining buffer (HBSS w/o calcium or magnesium, with 1% BSA), pelleted, split 50:50, and resuspended in ice-chilled FACS staining buffer (50 µL) and stained for either C3c (1:100-1:200 dilution) or C4d (1:200 dilution) on ice for 30 min in the dark. Cells were washed twice with FACS staining buffer (250-300 µL each wash) and either kept on ice (for C3c) or stained with secondary antibody (for C4d) on ice for 30 min (1:1000 dilution) in 50 µL. Secondary stained samples were washed two additional times, resuspended in ~280 µL FACS buffer and kept on ice until analysis. Samples were run on a Beckman Coulter Cytoflex S flow cytometer. After gating on single cells, a minimum of 10,000 events were collected for each sample. All cytometry data analysis was performed in Flow Jo (V10). Cells were evaluated for percent positivity by setting a gate at the histogram upper limit of the stained, heat-inactivated healthy control sample with the least histogram shift for each marker compared to the FMO control. Median fluorescence intensity (MFI) was calculated for each sample by subtracting raw MFI for each sample from the MFI for that marker on its own heat inactivated control. MFI could not be evaluated in some HEK293 C4d stained samples due to inconsistent amounts of secondary antibody. Percent positivity was preferred for C4d given its role as a complement initiator and MFI for C3c given its role in amplification. The same healthy control (n=4) and CM-HUS (n=4 for WT, n=5 for *PIGA*KO and *CD46*KO) samples were used across all cell types.

IgG and IgM

Flow cytometry for IgG and IgM deposition was similar to that of C3c and C4d with some notable differences. Briefly, 200k wild type cells were treated with heat-inactivated serum for 45 min at 37 °C. The only additional condition included DTT treatment of heat-inactivated serum (3 mM DTT pretreatment for

30 min at RT, 0.6 mM final concentration) of two CM-HUS samples. Cells were not split, but rather dual stained with AF647- conjugated goat anti-Human IgG or AF488-conjugated goat anti-Human IgM (Heavy chain) at a dilution of 1:100 for 30 min on ice. of IgG (HC n=9, CM-HUS n=17) or IgM (HC n=7, CM-HUS n=15). Relative median fluorescence intensity was calculated as ratio of sample MFI compared to average of 4 healthy controls ran with each experiment. *P* values were calculated using unpaired, two-tailed t test with Welch's correction.

**Supplemental Figures and Tables**

a

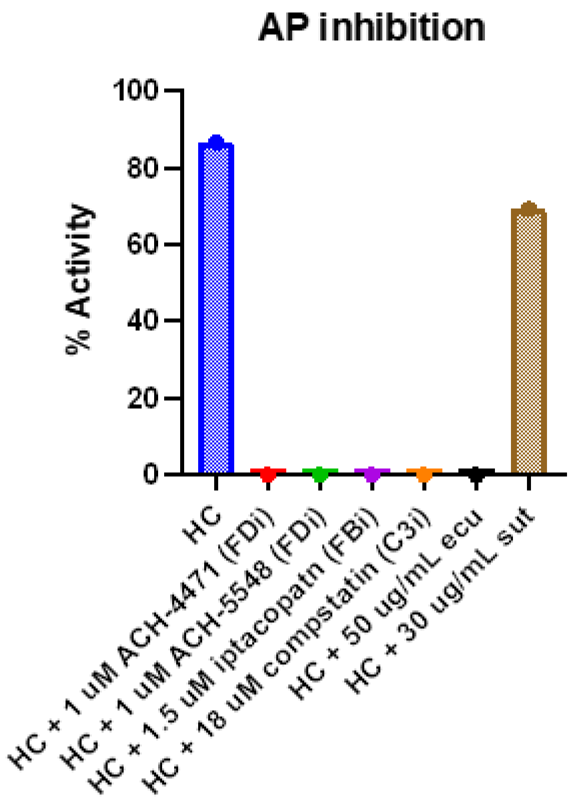


b

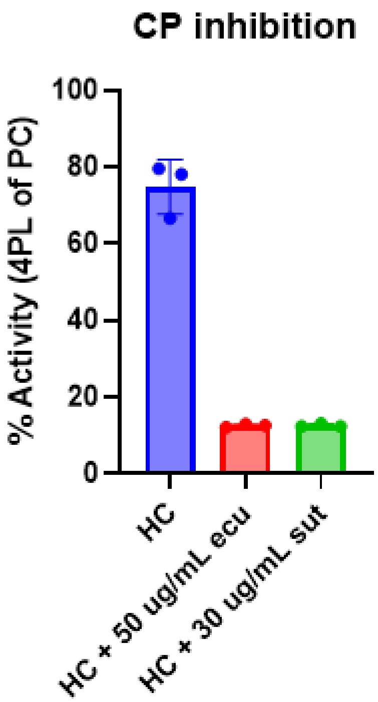


**Supplemental Fig. 1. a**, Wieslab AP inhibition. **b**, Wieslab CP inhibition. Each assay was completed as per manufacturer’s instructions using healthy control serum with or without inhibitors as documented.

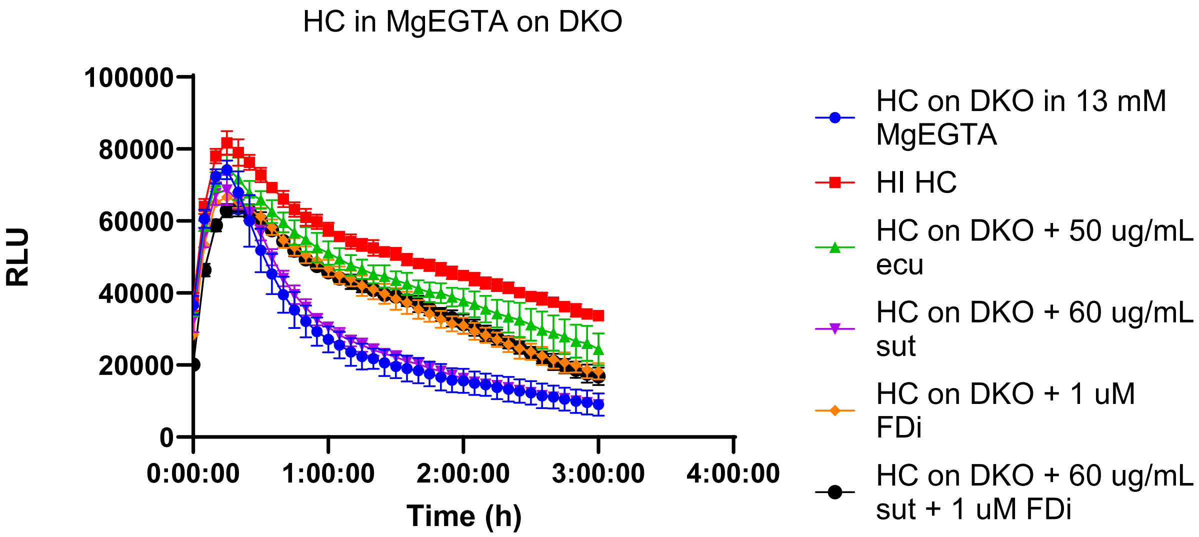


**Supplemental Fig. 2.** Assay in AP only buffer (13 mM MgEGTA in $GVB^0$) with healthy control serum on DKO cells. DKO cells were resuspended in MgEGTA with subsequent addition of healthy control serum with or without the addition of eculizumab, sutimlimab, Factor D inhibitor (FDi, ACH-5548), or a combination of sutimlimab and factor D inhibitor.

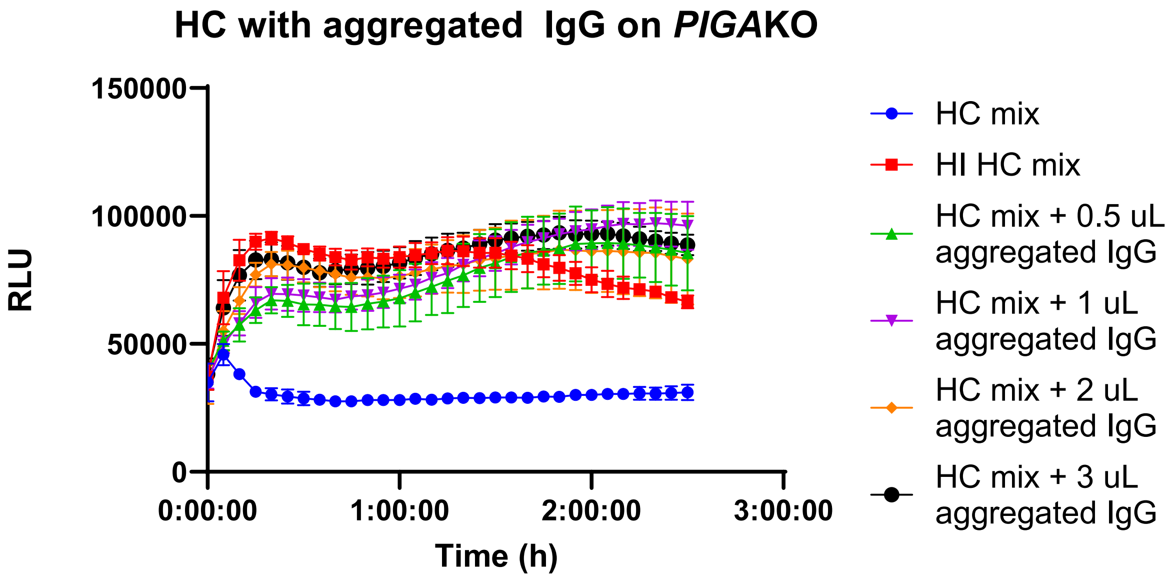


**Supplemental Fig. 3.** Heat aggregated IgG treatment of PIGAKO cells. Heat aggregated gamma globulin from Quidel (a strong CP stimulant) was added into the assay at the indicated volumes. Aggregated IgG was added to the cells to avoid complement depletion prior to running the assay.

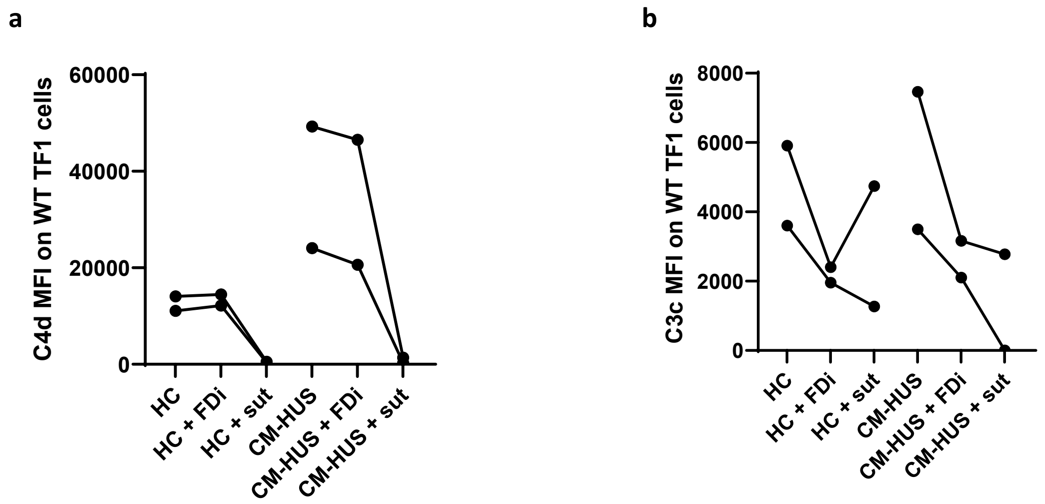


**Supplemental Fig. 4.** **a**,**b**, C4d (**a**) and C3c (**b**) median fluorescence intensity (MFI) on WT TF-1 cells with or without addition of sutimlimab (30 µg/mL, sut) or ACH-5548 (1 µM, FDi). HC (n=2) and CM-HUS samples (n=2) utilized are from same sera pools used on HEK293 cells. CR1 expression on TF-1 cells results in slightly different proportions of degradation fragments, ie better overall control of C3 in particular comparing HC to CM-HUS.

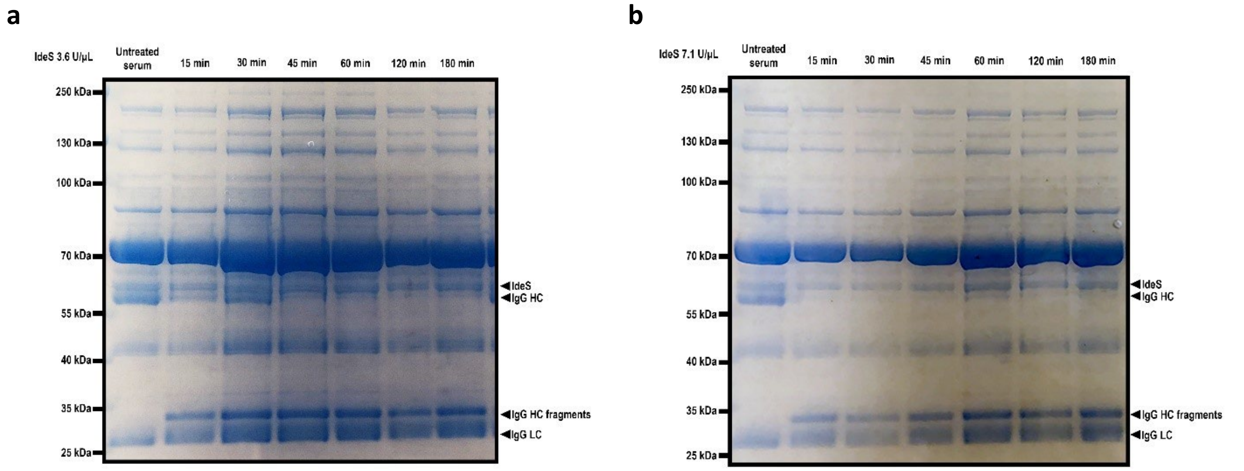


**Supplemental Fig. 5.** Optimized IdeS cleavage. NHS (10.4 µL of a 1:50 dilution) of NHS after treatment with IdeS at indicated timepoints. **a**, 3.6 U/µL IdeS. **b**, 7.1 U/µL IdeS.

**Supplemental Table 1.** Fractionated anti-sera sensitized sheep erythrocytes CH50.

| | Day 1 CH50 (U/mL) | Day 2 CH50 (U/mL) | Average CH50 (U/mL) |
|---|---|---|---|
| Untreated serum | 1300 | 1830 | 1565 |
| 0.6 mM serum pretreatment (0.006 mM final concentration) | 1070 | 1260 | 1165 |
| 3 mM serum pretreatment (0.03 mM final concentration) | 716 | 1350 | 1033 |
| 0.6 mM DTT final concentration | 953 | 772 | 862 |
| 3 mM DTT final concentration | 121 | 0 | 60.5 |

**Supplemental Table 2: Additional patient samples.**

| Patient identification | Age at diagnosis (y)/ Sex | Sample collected/ Follow-up (y) | Variant | Overall classification | Trigger | Therapy | Clinical outcome |
|---|---|---|---|---|---|---|---|
| C3G | 14/M | 2/10 | Heterozygous *CD46 c.38C>T, p.Ser13Phe* | VUS | NA | NA | ESRD awaiting transplant |
| HELLP01 | 33/F | 0/3 | NA | NA | Pregnancy | Supportive care only | CKD2 |

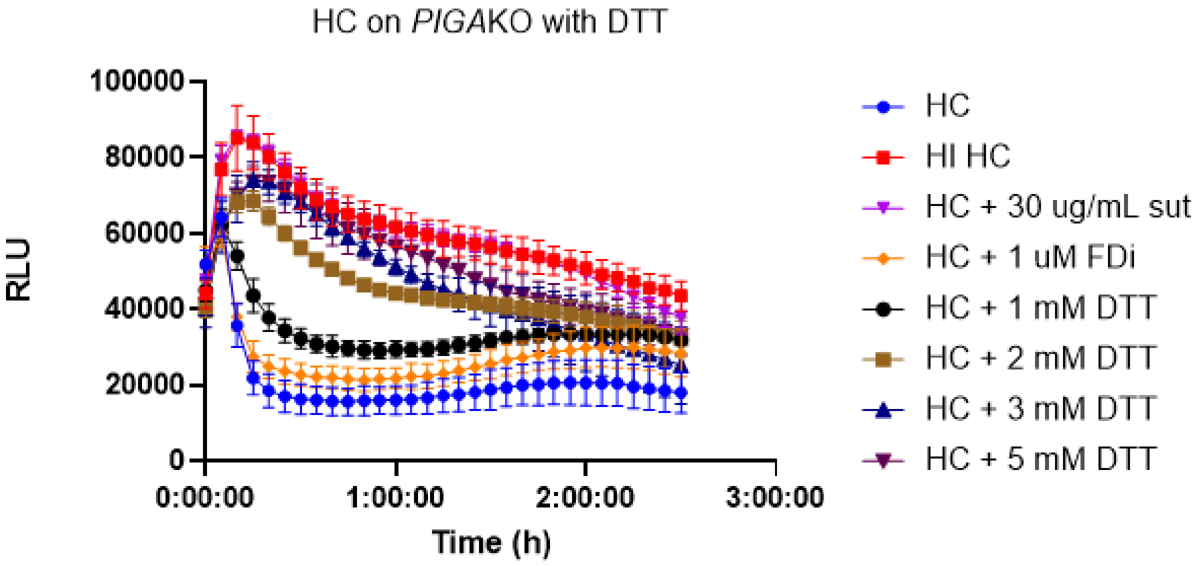


**Supplemental Fig. 6.** DTT treatment of HC sera on *PIGA*KO. HC sera is treated with increasing concentrations of DTT. Sutimlimab inhibition included for comparison.

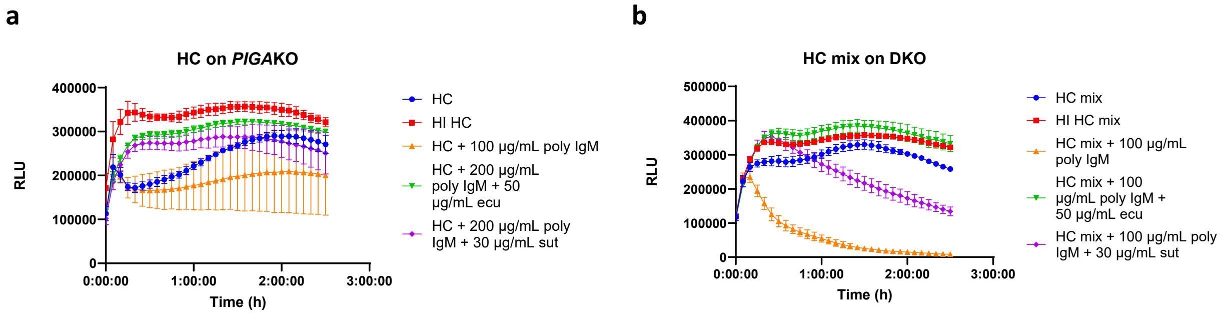


**Supplemental Fig. 7.** Inhibition of polyclonal IgM with eculizumab and sutimlimab on PIGAKO (**a**) or DKO (**b**). The data complements uninhibited data in Figure 5h,i.

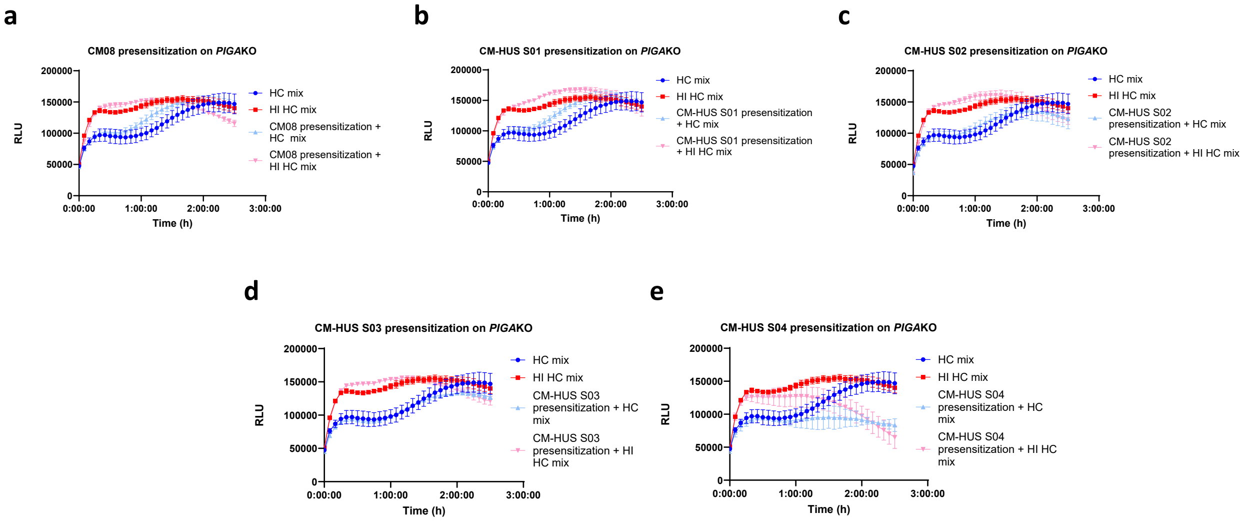


**Supplemental Fig. 8.** Presensitization of *PIGA*KO with CM-HUS serum in remission and or on C5 inhibition. **a-e,** CM-HUS-S01 through CM-HUS S04 are four CM-HUS patients in remission known to be on C5 inhibitor therapy not presented elsewhere in the manuscript. Specially selected low activity healthy control sera mix was utilized to facilitate complement activity in the bioluminescent mHam after PIGAKO cells were pre-sensitized (20% sera treatment for 45 min in DMEM), washed, and resuspended in GVB++ then ran in the bioluminescent mHam. Baseline activity of the healthy control mix sera shown in each tracing as dark blue (HC sera mix) and dark red (HI HC sera mix). Example traces plotted as mean +/- SD for each triplicate. RLU = relative luminescence units.
